# Aberrant NOTUM+ Program Induced in LGR5+ Crypt Base Columnar Cells Maintains an Immunosuppressive Niche in Colorectal Cancer

**DOI:** 10.1101/2025.03.10.642301

**Authors:** Julian Chua, Parnika Sakthivel, Arshdeep Kaur, Elleine Allapitan, Areeba Maqsood, Oliver F Bathe, Parham Minoo, Arshad Ayyaz

**Affiliations:** Department of Biological Sciences, University of Calgary, Calgary, Alberta, Canada; The Riddell Centre for Cancer Immunotherapy, Arnie Charbonneau Cancer Institute, Calgary, Alberta, Canada; Department of Oncology, Cumming School of Medicine, University of Calgary, Alberta, Canada; Department of Surgery, Cumming School of Medicine, University of Calgary, Alberta, Canada; Department of Pathology, Cumming School of Medicine, and Alberta Precision Laboratories, University of Calgary, Calgary, Alberta, Canada; The Calvin, Phoebe and Joan Snyder Institute for Chronic Diseases, Calgary, Alberta, Canada; Alberta Children’s Hospital Research Institute, Calgary, Alberta, Canada

## Abstract

Colorectal cancer (CRC) remains a leading cause of cancer-related mortality, with treatment failure largely driven by cancer stem-like cells that resist conventional chemoradiation and subsequently initiate tumor recurrence. While immune checkpoint blockade is effective in microsatellite instability-high (MSI-H) CRCs, the majority of CRCs are microsatellite stable (MSS) and exhibit immune exclusion, rendering them refractory to immunotherapy. Here, we identify a previously uncharacterized cancer cell subtype, which we term cancerous Crypt Base Columnar (canCBC) cells. These cells transcriptionally resemble normal LGR5+ CBC cells but activate an aberrant WNT/β-catenin signalling inhibitory program, marked by NOTUM expression. We show that canCBC cells are specifically enriched in MSS tumors, where their presence correlates with reduced CD8⁺ T cell infiltration, broader immune exclusion, and a propensity for regional lymphatic dissemination. Consistently, targeted ablation of canCBCs enhances the tumor-clearing potential of CD8⁺ T cells. This study identifies a novel therapeutic target for overcoming immune exclusion and improving immunotherapy responses in MSS CRCs.

---

Colorectal cancer (CRC) remains a leading cause of cancer-related mortality worldwide, with advanced-stage CRCs exhibiting high resistance to conventional therapies ^1,2^. While early-stage CRC is often treated effectively with surgery, patients with advanced disease require chemoradiation therapy and targeted treatments, which frequently fail to achieve long-term tumor control ^3,4^. A major barrier to successful treatment is the presence of cancer stem-like cells (CSCs), which survive therapies, reinitiate tumor growth, and drive metastatic progression ^5–8^. These therapy-resistant CSCs have long been known to exhibit quiescent or slow-cycling phenotypes, enabling them to evade chemotherapy, radiotherapy, and targeted agents designed to eliminate rapidly proliferating tumor cells ^9–11^. However, recent studies have demonstrated that the CSC compartment in CRCs is highly plastic and that radioresistant ‘revival’ CSCs (revCSCs), upon surviving therapeutic stress, can reconstitute a fresh pool of ‘proliferative’ CSCs (proCSCs), which rapidly expand, driving cancer reinitiation and progression, ultimately contributing to tumor recurrence and poor survival outcomes in CRC patients ^12,13^.

Advances in immunotherapy, particularly immune checkpoint blockade (ICB), have revolutionized the treatment of many cancers, especially microsatellite instability-high (MSI-H) CRCs that exhibit high mutation burdens and strong immune infiltration ^14,15^. However, most CRCs (∼85%) are microsatellite stable (MSS) and remain refractory to ICB therapies, likely due to their immune-excluded tumor microenvironment ^16,17^. Furthermore, immunotherapies can induce severe immune-related toxicities, limiting their widespread application in CRC patients ^18,19^. Despite numerous efforts to improve treatment responses, a major gap remains in understanding the molecular programs that underlie immune evasion in MSS CRCs, particularly in the context of CSC populations that have now been shown to promote resistance against immunotherapies as well ^20–22^.

Here, we performed a comprehensive single-cell transcriptional analysis of CRC tissues across five independent patient cohorts to investigate the heterogeneity of cancer stem-like populations and their relationship with immune exclusion. Our analysis identified a previously uncharacterized cell population, which we term cancerous Crypt Base Columnar (canCBC) cells due to their transcriptional resemblance to normal LGR5+ CBCs but with an aberrant, tumor-associated transcriptional program that is absent from normal colonic epithelium. This program is characterized by elevated expression of multiple WNT/β-catenin signalling inhibitors, including NOTUM, NKD1, and APCDD1, suggesting a key role in tumor progression and immune modulation. Notably, canCBCs are specifically enriched in MSS CRCs, where they correlate with immune exclusion, particularly reduced CD8⁺ T cell infiltration, and lymph node metastasis. Consistent with these observations, targeted ablation of canCBC cells promotes the ability of activated CD8+ T cells to remove cancerous cells. These findings indicate that canCBCs significantly contribute to the immune-resistant phenotype of MSS CRCs, potentially serving as a novel therapeutic target for overcoming immune evasion and treatment resistance.

## Results

### Deciphering heterogeneity in cancer stem-like cells

We analysed transcriptomes from 639,249 single cells originally isolated from primary CRC tissues across 140 patients, encompassing five independent patient cohorts accessible through public sources ^23–27^. To ensure robust cross-dataset alignment while preserving biological variability, we utilized Harmony, a batch correction and data integration algorithm optimized for large-scale single-cell transcriptomic datasets ^28^. This approach effectively mitigated technical variability, aligned shared cellular states across datasets, thereby enhancing the comparative analysis of distinct CRC patient cohorts. This was evident in the clear separation of major cellular compartments, including epithelial (both malignant and normal colonic), lymphoid (T cells, B cells, plasma cells), myeloid (macrophages, monocytes, dendritic cells), mesenchymal, and endothelial populations, across all five patient cohorts (**Extended Data Fig. 1a,b**).To investigate cellular heterogeneity within the cancerous tissues, we isolated cancerous single cell transcriptomes and performed unsupervised clustering. Using previously established gene signatures ^13,29^, we identified distinct subpopulations, including proCSCs (two populations were detected mainly distinguished by variations in rate of proliferation), revCSCs, and differentiated absorptive and secretory lineages (**Fig. 1a**). Consistent with previous studies, both proCSC subtypes represented the most proliferative cell populations in human CRC tissues. The proCSC1 subtype exhibits the highest proliferation rate, with over 95% of cells in the S, G2, or M phase. In contrast, proCSC2 displays a lower – but still higher than any non-proCSC subtype proliferative index, with 28% of cells in S/G2/M phases (**Fig. 2b**). In contrast, revCSCs displayed a quiescent or slow-cycling phenotype, with 86% of cells arrested in G1 or G0 phase. Similarly, the vast majority of secretory (81%) and absorptive (89%) lineages were predominantly in G1 or G0, indicative of their differentiated state. Our analysis also revealed a previously uncharacterized transcriptional subtype (cluster #6) that exhibited a relatively quiescent phenotype (79% in G1 phase, 14% in S, and 8% in G2/M). These cells were distinguished by the ectopic expression of multiple WNT/β-catenin signalling inhibitors, including NOTUM, NKD1, and APCDD1 (**Fig. 1a**). Together, these results show that the stem cell-like compartment in human CRCs is heterogenous.

**Figure 1.**
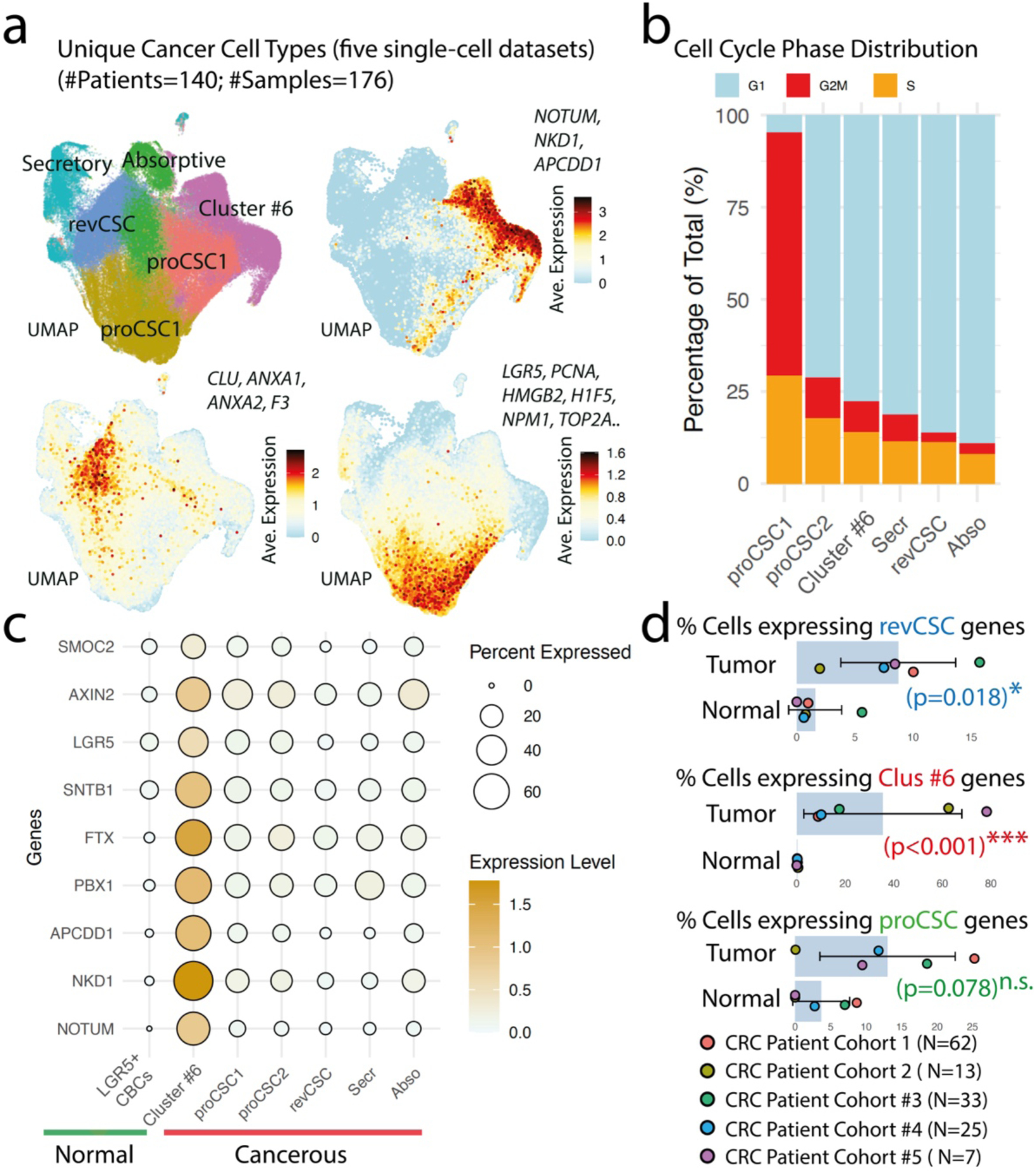
Aberrant NOTUM+ Transcriptional Program is Cancer Specific. **a**, A UMAP visualization of single-cell transcriptomes of cancerous cells from five independent CRC patient cohorts, with average gene expression patterns shown for markers of three identified cancer cell subtype, proCSC, revCSC and unknown cluster #6. **b**, Cell cycle distribution analysis across all identified cancer cell subtypes confirming that proCSCs are most proliferative, revCSCs are least proliferative while cells in cluster #6 are moderately proliferation CSC subtypes. **c**, Dotplot is showing expression levels of indicates genes in all identified cancer cell subtypes in comparison with normal LGR5+ crypt based columnar (CBC) cells detected in matching adjacent normal colonic epithelia. **d**, The percent abundance of cells expressing gene signatures of revCSCs (ANXA1, CLU, ANXA2, F3), cluster #6 (NOTUM, APCDD1, NKD1), and proCSCs (HMGB2, TUBA1B, PCLAF, UBE2C, DUT, TOP2A, H1F5, HSPD1, NCL, BIRC5, NPM1, RANBP1, CDK1, HSPE1, RAN, PCNA, GMNN, RRM2, H2AZ1, DTYMK, SNRPD1, LYAR, LSM2, NHP2) in normal and cancerous tissues was calculated for each of the five CRC patient cohorts and compared for statistical significance using a paired Student’s t-test; n.s., non-significant. ‘N’ represents the number of CRC patients in each cohort.

**Figure 2.**
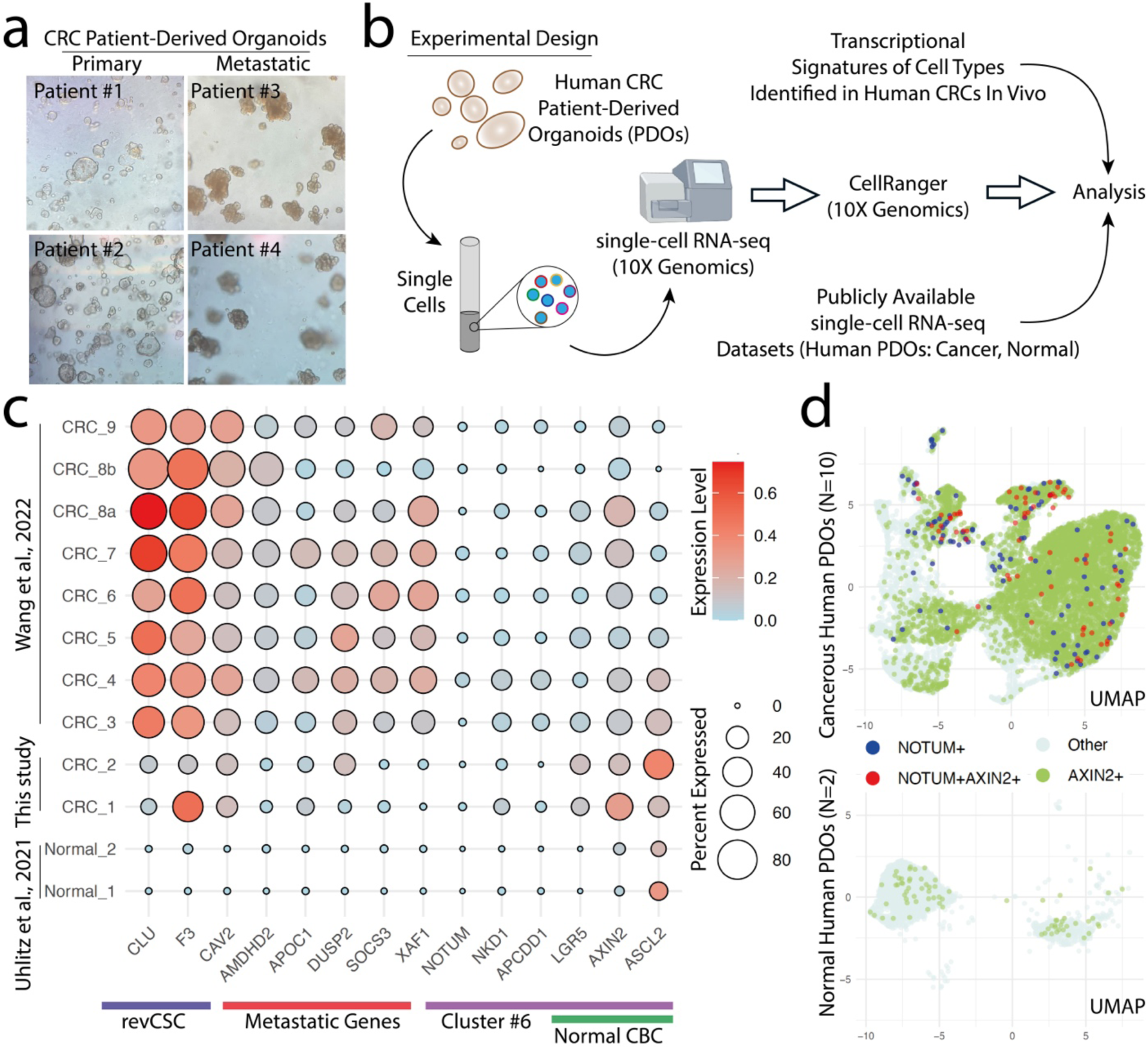
Colorectal Cancer Patient-Derived Organoids Sustain NOTUM+ Program Activation. **a**, Representative images of four patient-derived organoids (PDOs). **b**, Schematic of experimental design and subsequent analysis. **c**, Expression patterns, including expression levels and the percentage of cells expressing the indicated genes, are shown in cancerous human PDOs and compared with normal human organoids, confirming that the revCSC and aberrant NOTUM⁺ transcriptional programs are retained exclusively in cancerous organoids and remain absent in normal colonic organoids. Note that single-cell transcriptomes of CRC_1 and CRC_2 generated in this study (which correspond to PDOs of Patient #1 and Patient #2 shown in panel (**a**), respectively) were resected from primary tumors in the colon. **d**, A UMAP representation of single cells expressing NOTUM (a specific cluster #6 marker) and AXIN2 (expressed in cluster #6 and previously shown to mark normal LGR5⁺ CBC cells as well), color-coded and labeled as positive when the normalized, log-transformed expression value of each gene was greater than zero.

### LGR5+ cancer cells induce a distinct NOTUM+ transcriptional program absent from normal gut epithelia

Prior studies have shown that a cancer-associated transcriptional program similar to the one we observed in cluster #6 emerges in LGR5⁺ CBC cells specifically at cancer onset, where it was thought to confer a competitive advantage, facilitating clonal expansion within the crypt niche ^30,31^. Indeed, cluster #6 also exhibited the high expression of LGR5 and several other canonical CBC markers, such as AXIN2 and SMOC2, although their basal expression across all cancer-associated clusters was relatively higher when compared with normal CBC cells (**Fig. 1c**). Moreover, the transcriptional program uniquely activated in cluster #6, which included NOTUM, NKD1, APCDD1, PBX1, FTX1 and SNTB1, was mostly absent in normal LGR5⁺ CBCs from matching adjacent normal colonic tissues of CRC patients across all five patient cohorts (**Fig. 1c**, **Extended Data Fig. 2a,b**).

To evaluate the robustness of these findings, we quantified the proportion of cancer cells expressing the most significantly enriched cluster #6-specific genes (NOTUM, NKD1, and APCDD1) that were absent from normal LGR5+ CBC cells, and compared them to other CSC subtypes, classified based on their respective gene signatures across five patient cohorts. Using paired statistical tests, we assessed the relative proportions of these CSC populations within each patient cohort and evaluated whether the observed trends were significant across all five cohorts. Our analysis revealed that transcriptional programs active in both cluster #6 cells and revCSCs were exclusively and significantly upregulated in cancerous tissues (**Fig. 1d**). In contrast, the proCSC program, representing the most rapidly proliferating cells in human CRC, was paradoxically abundant in both tumor and normal tissues, showing no significant difference between them across five patient cohorts.

Together, our findings reveal two key CSC phenotypes – cluster #6 cells and revCSCs – whose transcriptional programs are specifically associated with cancerous tissues.

### NOTUM⁺ program represents an inherent property of cancerous LGR5⁺ cells

CRC cells exist within a complex microenvironment that significantly influences their transcriptional responses and behavior. To determine whether the CRC phenotypes we observed were intrinsic to cancerous tissues rather than induced by their surroundings in vivo, we established patient-derived organoids (PDOs) from four primary CRC resection specimens: two from primary tumors and two from metastatic liver lesions of invasive cancers (**Fig. 2a**). Stable PDO cultures from primary tumors were then subjected to single-cell RNA sequencing (scRNA-seq), and results were compared with publicly available single-cell transcriptomes from eight additional CRC PDOs and two normal colonic samples ^26,32^ (**Fig. 2b**). Consistent with our in vivo findings, revCSC markers and several key genes associated with cluster #6 cells were differentially induced in multiple cancerous PDOs compared to organoids derived from normal colonic epithelia (**Fig. 2c**). Additionally, the expression levels of canonical CBC markers, including LGR5, AXIN2, and ASCL2, were significantly higher in cancerous PDOs compared to their basal levels in normal organoids. Furthermore, we observed that the cluster #6 signature was predominantly expressed in PDOs that also expressed genes associated with metastatic invasion, whereas the revCSC-specific gene expression pattern showed no correlation with these genes ^33^. Notably, no LGR5 transcripts were detected in normal organoids, potentially due to low basal expression levels that were undetectable through single-cell profiling technique applied in these assays. To further validate our findings, we therefore employed another highly specific CBC cell marker AXIN2 ^34^, and labeled AXIN2⁺ (single-positive), NOTUM⁺ (single-positive) and AXIN2⁺ NOTUM⁺ (double-positive) cells, defining positive labeling as normalized, log-transformed expression values greater than zero. Indeed, while a large number of NOTUM⁺ (single-positive) and AXIN2⁺ NOTUM⁺ (double-positive) cells were detectable in cancerous PDOs, not a single NOTUM⁺ or AXIN2⁺ NOTUM⁺ double-positive cell was detected in normal organoids (**Fig. 2d**). Additionally, a significantly higher number of AXIN2⁺ (single-positive) cells were observed in cancerous PDOs, whereas only a few were present in normal organoids.

Together, these findings show that the constitutive activation of NOTUM⁺ transcriptional program is distinct, represents an intrinsic feature of cancerous cells rather than an adaptation to their microenvironment and is generally inactive in normal epithelial regions of the human gut. Given these observations, and because cluster #6 also express canonical CBC markers, we designate these cells as cancerous CBC (canCBC) cells.

### Enrichment of canCBC program is inversely associated with CRC patient survival

To assess the clinical relevance of the identified CSC subtypes, we applied the Gene Set Variation Analysis (GSVA) method ^35^ to stratify CRC patients from the TCGA-COAD dataset based on subtype-specific gene signatures. A subset of 427 patients who had undergone conventional chemoradiation therapy with recorded clinical follow-up data was selected for this analysis. Unbiased clustering revealed that proCSC-enriched patient samples clustered with those exhibiting replicating cell gene signature, consistent with their highly proliferative phenotype, thereby validating the utility of applying single-cell transcriptional phenotypes to bulk RNA-seq for patient stratification. Notably, revCSC- and secretory cell-enriched patients were closely associated, suggesting a potentially transcriptional overlap between these two subtypes (**Fig. 3a**). In contrast, absorptive and canCBC-enriched patients formed distinct clusters, indicating transcriptional divergence from other tumor subtypes.

**Figure 3.**
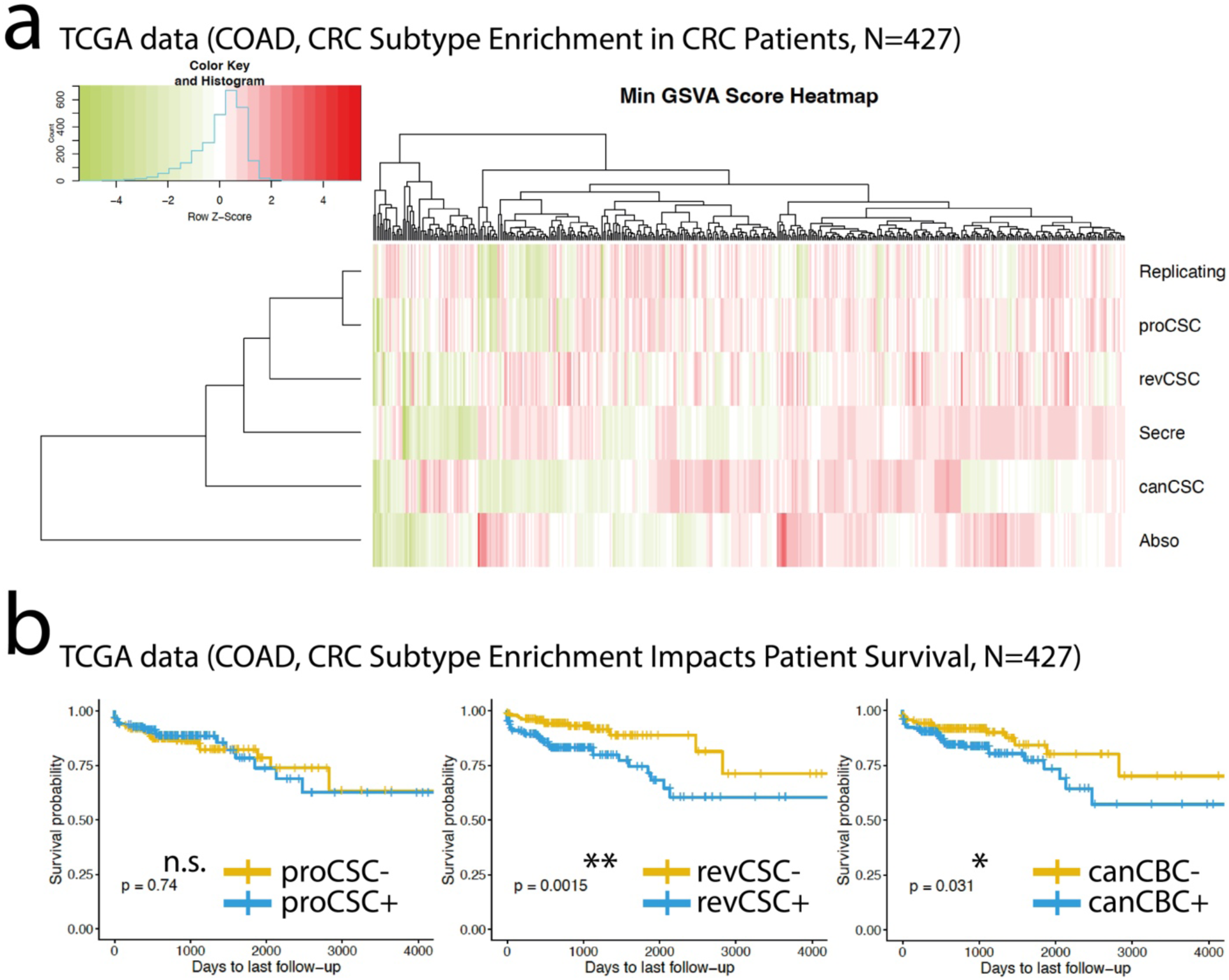
Enrichment of canCBC Program Associates with Poor Clinical Outcome in Colorectal Cancer Patients. **a**, Enrichment scores of indicated transcriptional programs were calculated for 427 patients individual and plotted on a heatmap. **b**, Kaplan-Meier (KM) survival curves for patients enriched (+) or deficient (-) in the indicated transcriptional programs. Significance was tested using the log-rank test, which calculates p-values based on the χ² (chi-square) statistic.

Next, we used CRC subtype enrichment scores to classify patients into binary subcohorts, defining them as either enriched (‘+’) or deficient (‘-’) for each CSC-specific transcriptional program. This stratification revealed that enrichment of both revCSC- and canCBC-specific transcriptional programs was significantly associated with worse survival outcomes in CRC patients undergoing chemoradiation therapy, whereas proCSC enrichment had no significant impact on patient survival (**Fig. 3b**).

Together, our results show that revCSC and canCBC subtypes contribute to decreased survival in CRC patients undergoing chemoradiation therapies.

### NOTUM+ canCBC program is selectively induced in regionally disseminating MSS type colon cancers

Advanced CRCs frequently develop resistance to conventional chemoradiation therapy, suggesting that revCSC and canCBC cells may survive clinical treatments and drive tumor recurrence due to their stem cell-like properties, ultimately leading to poor clinical outcomes ^2,36^. However, another critical determinant of therapy response is the extent of immune cell infiltration into the tumor microenvironment, which in human CRCs can be largely predicted by the microsatellite status – repetitive DNA sequences prone to mismatch repair deficiencies ^37^. This parameter also determines the efficacy of more recently developed anti-CRC immunotherapies ^38^. Thus, microsatellite instability-high (MSI-H) tumors are characterized by high mutation burdens, rendering them highly immunogenic and responsive to immune checkpoint blockade therapies ^2,38–40^. In contrast, microsatellite stable (MSS) tumors exhibit low mutational loads and are generally refractory to immunotherapy. Fortunately, one of the five datasets analysed included MSI-H/MSS classification data from 62 CRC patients ^23^. This allowed us to examine the relative distribution of canCBC, revCSC, and proCSC cells across tumor subtypes. Our analysis revealed that canCBC cells were exclusively enriched in MSS tumors, whereas revCSCs were relatively more abundant in MSI-H tumors, while proCSCs were distributed equally across both subtypes (**Fig. 4a,b**). In addition, secretory and absorptive gene signature expressing cancer subtypes were also equally distributed among MSS and MSI-H CRCs.

**Figure 4.**
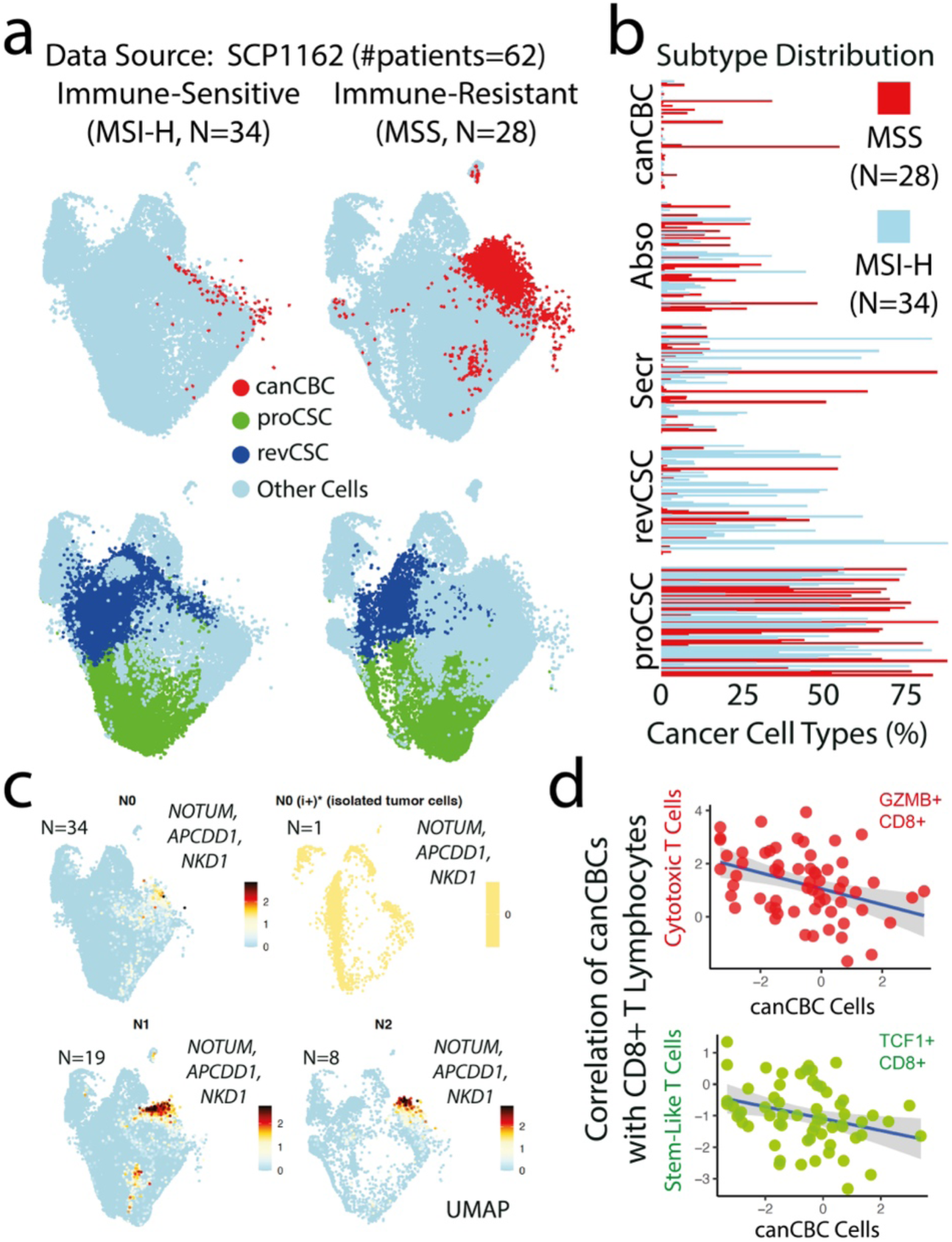
canCBC Program Activation in Microsatellite Stable Colorectal Cancers. **a**, UMAPs showing that canCBCs (red dots) are predominantly found in MSS, but not MSI-H, CRC cases. It is worth noting that proCSCs (green dots) are prevalent in both MSS and MSI-H CRCs, while revCSCs (blue dots) are mostly abundant in MSI-H CRC cases. **b**, Percent distribution of all identified CRC cell subtypes is calculated among individual MSS and MSI-H cases. **c**, Correlation analyses between the frequencies of NOTUM+ and stem-like/cytotoxic T cells. **d**, Correlation analyses between the frequencies of NOTUM+ and stem-like/cytotoxic T cells. N represents number of CRC patients represented in the corresponding analysis. ‘N’ represents the number of CRC patients.

We also observed that only a subset of MSS CRCs contained a substantial number of canCBC cells, prompting a closer examination of the clinical information available for the various patient cohorts (**Fig. 4b**). Interestingly, while there was no direct correlation between the enrichment of canCBC or proCSC gene signatures and tumor size, a higher proportion of revCSCs was directly associated with larger tumors (**Extended Data Fig. 3**). Next, we examined a potential link between these CSC programs and pathological tumor (pT) staging. This revealed that while revCSC- and proCSC-specific transcription was abundant in all pT stages, pT4 cases exhibited the highest levels of the canCBC gene signature, followed by pT3, while only a few cells in pT2 and pT1 cases expressed canCBC signature (**Extended Data Fig. 4**). This enrichment of canCBC program in pT4/pT3 tumors suggested that these cancers have potentially metastasized into adjacent lymph nodes. To assess whether canCBC cells associate with regional lymphatic dissemination, we examined their distribution across nodal stages and found that canCBC cells were also enriched in N1 and N2 tumors but absent in N0 (**Fig. 4c**). In contrast, revCSC and proCSC programs were indiscriminately expressed across all nodal stages (**Extended Data Fig. 5**).

Clinical observations have linked lymph node metastasis with the deficiency of cytotoxic T cells ^41^. Therefore, to explore the relationship between cytotoxic T cell infiltration and canCBC enrichment, we quantified CD8⁺ T lymphocyte infiltration in tumors, distinguishing between cytotoxic CD8⁺ T cells (co-expressing GZMB and CD8A) and stem-like CD8⁺ T cells (co-expressing TCF7 and CD8A). We then correlated their infiltration levels with canCBC cell abundance across CRC patients. This analysis revealed a negative correlation between CD8⁺ T cell infiltration and canCBC cell enrichment, suggesting that tumors harbouring high canCBC populations exhibit an immune-excluded phenotype (**Fig. 4d**).

Together, these results show that the canCBC program is specifically activated in MSS CRCs, progressively upregulated as cancerous tissues disseminate into nearby tissues and lymph nodes and negatively associated with CD8⁺ T cell count.

### NOTUM+ canCBC program associates with immune cell exclusion

To determine whether the lack of immune cell infiltration was specific to CD8⁺ T cells or reflected a broader immune desert phenotype, we turned to the TCGA-COAD dataset, which we had previously analysed (see **Fig. 3**). Using Gene Set Variation Analysis (GSVA) ^35^, we assigned immune cell-type enrichment scores based on validated immune signature gene sets from prior studies ^35^. Strikingly, among all identified CRC subtypes, canCBC cells exhibited the lowest association with the immune cell types (**Fig. 5a**). In addition, while revCSC and proCSC cells were positively associated with multiple immune populations, including activated CD8⁺ and CD4⁺ T cells, canCBC cells were negatively correlated with most immune cell subsets, including activated CD8⁺ and CD4⁺ T cells. One notable exception was a positive correlation between canCBC cells and CD56⁺ bright natural killer (NK) cells, a subset primarily involved in antimicrobial immunity rather than tumor immunosurveillance, prompting us not to further pursue this observation in the context of current study.

**Figure 5.**
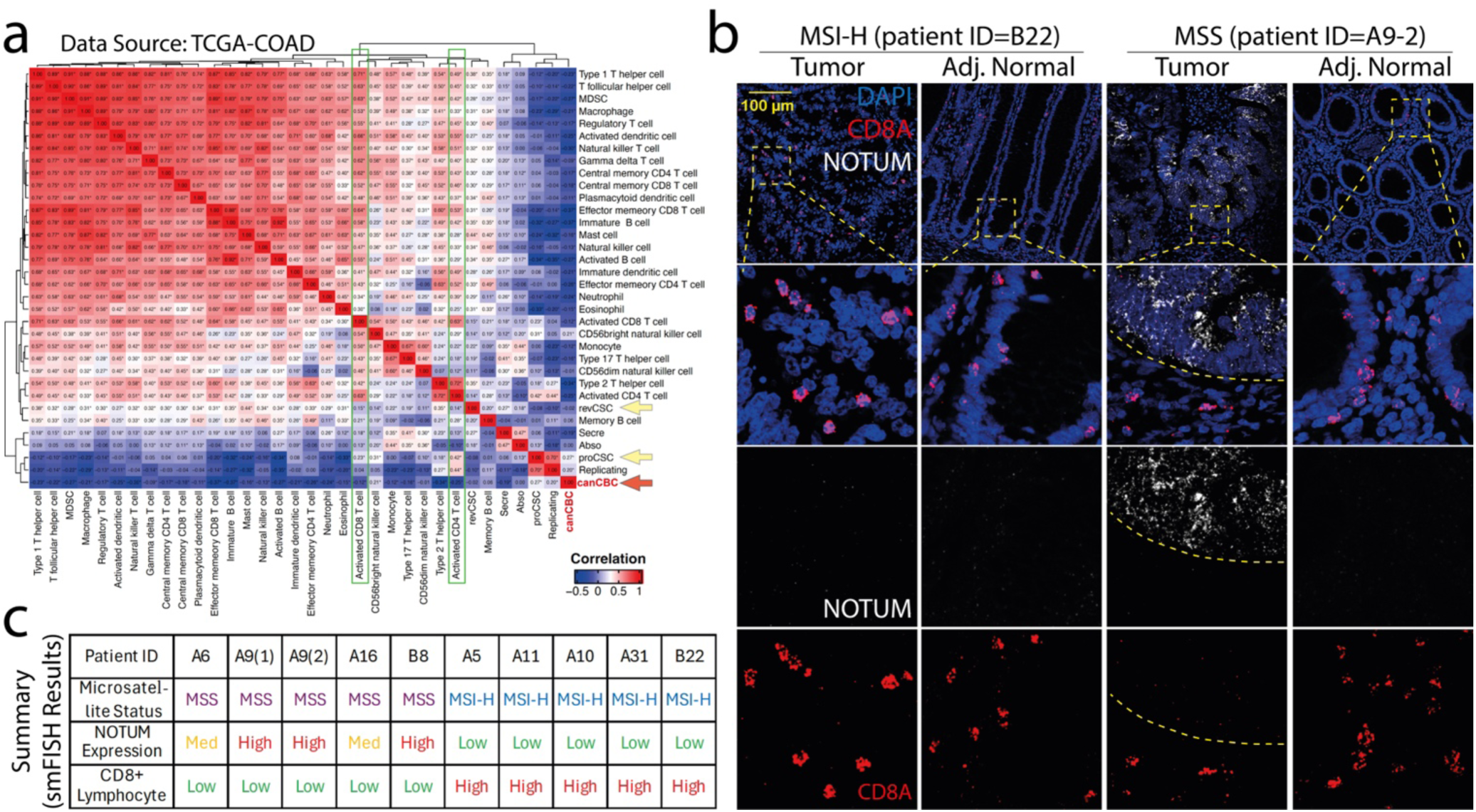
canCBC Program Associates with Immune Exclusion. **a**, Heatmap showing correlations between indicated cell types, where the color bar represents the correlation coefficient while the statistical significance is shown as: *: p<0.05, **: p<0.01, ***: p<0.001. Note that canCBCs (indicated with green thick arrow; while revCSCs and proCSCs are indicated with yellow thick arrows) are least associated with the abundance of immune cells in the tumor in general. **b**,**c**, Representative images (**b**) from the RNAScope-based single-molecule in situ hybridization (smFISH) analysis of two primary CRC patient samples showing that NOTUM is specifically expressed in MSS tumors, and is linked to lack of CD8+ T cell infiltration. Note that CD8+ T cells invade and survey MSI-H tumors as well as normal colonic epithelia. These results from all ten samples are summarised in (**c**).

To independently validate these findings, we examined fresh tissue samples from a CRC patient cohort unrelated to the five cohorts analysed above. Case recruitment was based on two criteria: (i) MSS or MSI-H CRC scored as pT4, and (ii) all samples must be accompanied by matching adjacent normal gut epithelium for direct comparison within each patient sample. A total of ten resection specimens from nine patients were obtained, including four MSS and five MSI-H CRC cases. We then performed single-molecule in situ hybridization (smFISH) to assess the expression levels of NOTUM and CD8A genes (**Fig. 5b**). These analyses confirmed that NOTUM was selectively upregulated in cancer cells within MSS tumors but was absent in MSI-H tumors. Furthermore, NOTUM expression was strongly associated with lack of CD8⁺ T cell infiltration (**Fig. 5b,c**; representative smFISH images from one MSS and one MSI-H case illustrate these findings in (**b**), while summary of results from all analysed patient samples are presented in (**c**)). Importantly, NOTUM expression was not detected in adjacent normal colonic tissues from either MSS or MSI-H patients, underscoring its tumor-specific activation.

Consistent with our results from single-cell transcriptome analyses, CD8⁺ T cells exhibited extensive infiltration in MSI-H tumors, with CD8⁺ T cells actively surveying adjacent normal colonic epithelium in both MSS and MSI-H cases.

Together, these results show that canCBC enrichment is a hallmark of immune excluded human CRCs.

### Targeted ablation of canCBC cells enhances CD8⁺ T cell activation and eliminates colorectal cancer cells

To investigate whether canCBC cells directly suppress CD8⁺ T lymphocyte activity, we developed an adoptive cell therapy (ACT) model utilizing murine AKP organoids and OT-1 T cell co-culture system ^42^. These organoids were derived from normal intestinal organoid cultures established from C57BL/6 mice and engineered using CRISPR-Cas9 to delete Apc and p53 genes, while simultaneously introducing a previously designed KrasG12D-GFP transgene ^43^ (**Fig. 6a**). Efficiency of Apc and p53 knockout and successful incorporation of KrasG12D-GFP were confirmed by quantitative RT-PCR, comparing gene expression between AKP cancerous and matching normal organoids (**Extended Data Fig. 6a,b**).

**Figure 6.**
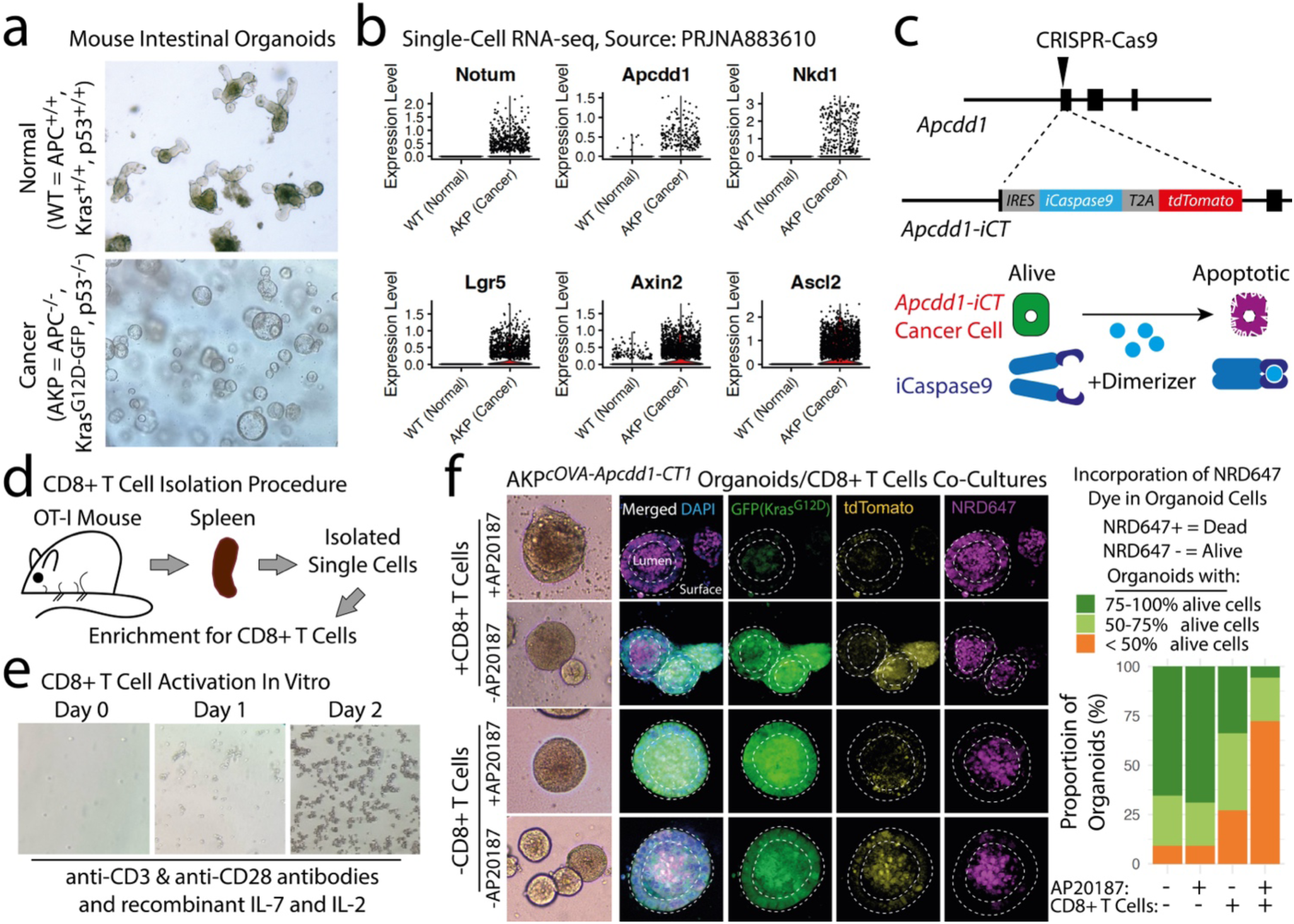
Targeted Ablation of canCBC Cells Induces CD8+ T Cell Mediated Removal of Cancer Cells. **a**, Representative images of normal and AKP (APC-/-, KrasG12D-GFP, p53-/-) mouse organoids is shown. **b**, Violin plots showing that AKP mouse organoids express high levels of CBC marker genes, such as Lgr5, Axin2 and Ascl2, as well as aberrant canCBC-specific genes, such as Notum, Apcdd1 and Nkd1. **c**,**d**, Schematics showing the strategy used in the generation of Apcdd1-iCT transgene (**c**), and extraction of CD8+ T cells from the spleens of OT-1mice. **e**, Images showing in vitro expansion of OT-1 CD8+ T cells at indicated timepoints following isolation and enrichment. **f**, Results from AKP tumor organoid-CD8+ T cell co-cultures are shown.

Since AKP organoids model immunosuppressive MSS colorectal tumors, as previously demonstrated ^44^, we first sought to determine whether they exhibit the canCBC transcriptional program. Single-cell RNA-seq analysis revealed a distinct canCBC-like gene expression signature, including upregulation of Notum, Apcdd1, and Nkd1, alongside higher expression of canonical CBC markers, such as Lgr5, Axin2, and Ascl2, compared to wild-type (WT) intestinal organoids (**Fig. 6b**, **Extended Data Fig. 7**). Thus, murine AKP organoids recapitulate canCBC-enriched MSS type human CRCs.

Previous studies suggest that Notum may also promote tumor growth in a manner independent of their interaction with the immune system ^45^. Therefore, to enable selective and inducible ablation of canCBC cells, we selected Apcdd1 gene, another highly specific canCBC marker, to develop a dual-function AKP-Apcdd1-iCT organoid model, which serves both as a tdTomato-based canCBC reporter and as a platform for targeted canCBC cell depletion. This system utilizes AP20187, a chemical inducer of dimerization (CID) that activates the FKBP fusion protein system, enabling spatiotemporally controlled elimination of canCBC cells, as previously demonstrated ^5^ (**Fig. 6c**). Furthermore, we engineered these organoids to express the chicken ovalbumin peptide SIINFEKL (cOVA), facilitating antigen-specific recognition by SIINFEKL-restricted CD8⁺ T cells (OT-I T cells). To generate activated CD8⁺ T cells, OT-I T cells were isolated from mouse spleens and stimulated ex vivo using anti-CD3 and anti-CD28 antibodies and inflammatory cytokines interleukin-7 (IL-7) and IL-2 (**Fig. 6d,e**). Next, AKP-cOVA-Apcdd1-iCT organoids were cultured in the presence or absence of activated OT-I T cells, followed by prior AP20187-mediated dimerization-induced canCBC cell ablation. As an additional control, AKP-cOVA-Apcdd1-iCT co-cultures with CD8⁺ T cells were maintained without dimerizer treatment. Organoids were analysed after 48 hours of co-culture.

Our results show that CD8⁺ T cells preferentially clustered around organoids that lacked canCBC cell population, with many of these organoids displaying structural collapse (**Fig. 6f**). This phenotype was not observed under control conditions, which includes organoids treated with AP20187 alone in the absence of CD8⁺ T cells; these organoids remained intact, indicating that tumors undergoing canCBC cell ablation are capable of fully recovering in the absence of cytotoxic T cells. Additionally, tumor organoids that were not treated with the AP20187 dimerizer prior to co-culture with cytotoxic T cells also remained structurally intact, suggesting that canCBC cells play a crucial role in maintaining organoid integrity following exposure to T cells.

To quantitatively examine apoptotic induction in organoid cells, we incubated the cultures for 30 minutes with NucRed Dead 647, a membrane-impermeable dye that selectively penetrates cells undergoing apoptosis and binds to DNA, emitting fluorescence in the far-red spectrum. Organoids were then scored based on the proportion of apoptotic cells, categorizing them into three groups according to the fraction of NucRed Dead 647-positive (Dead) and negative (Alive) cells (**Fig. 6f**). Our analysis demonstrated that AP20187-mediated canCBC cell ablation alone had no significant effect on organoid viability, confirming that targeted elimination of canCBC cells in the absence of CD8⁺ T cells does not compromise organoid integrity. This was further substantiated by the presence of abundant GFP⁺ cells, indicative of a KrasG12D-GFP transgene-expressing tumor cell population (**Fig. 6f**). In contrast, co-culture with activated OT-I CD8⁺ T cells alone, in the absence of canCBC ablation, as evident by clear tdTomato signal in these cells, resulted in moderate levels of tumor cell death but did not induce complete organoid collapse. Despite CD8⁺ T cell-mediated cytotoxicity, these tumor organoids remained structurally intact and retained GFP signal, suggesting that while cytotoxic T cells can recognize and attack some tumor cells even in the presence of canCBC cells in our in vitro co-culture model, the tumor organoids are able to withstand this immune pressure and avoid complete collapse and elimination.

However, in conditions where canCBC cells were selectively ablated prior to co-culture with cytotoxic T cells, we observed markedly enhanced tumor cell death, with 72% of organoids exhibiting more than 50% apoptotic cells (NucRed Dead 647-positive) (**Fig. 6f**, **Extended Data Figure 8a,b**). This was significantly higher than the 27% observed in cultures with CD8⁺ T cells alone and the 9% in organoids subjected to canCBC cell ablation without T cell exposure, a proportion that mirrored untreated controls, where only 9% of organoids contained less than 50% apoptotic cells.

Together, these findings demonstrate that targeted depletion of canCBC cells prior to exposure to cytotoxic T cells significantly enhances anti-tumor immune responses, leading to greater tumor cell elimination.

## Discussion

Despite significant advancements in CRC therapy, resistance to chemoradiation, immune evasion, and toxicity to healthy tissues remain major barriers to improving patient outcomes. A subset of CSCs has been implicated in tumor recurrence, metastasis, and therapy resistance, largely due to their ability to evade conventional treatments and repopulate tumors following therapeutic failure ^46^. However, many CSC populations share molecular signatures with normal stem cells, limiting their potential as therapeutic targets due to the risk of off-target toxicity and damage to normal tissues.

In contrast, the canCBC population identified in this study, while expressing high levels of intestinal stem cell markers, undergoes a cancer-specific transcriptional reprogramming that distinguishes it from normal LGR5⁺ CBC cells. This aberrant program creates a unique therapeutic vulnerability, allowing for selective targeting of canCBCs while sparing normal stem cells.

While ICB has demonstrated efficacy in MSI-H CRCs, confirming the feasibility of immunotherapy in colorectal malignancies, MSS CRCs (∼85% of cases) remain refractory to these treatments due to their immune-excluded tumor microenvironment ^15,38^. This study identifies canCBC cells as a previously uncharacterized CSC subset that emerges during tumor development and is enriched in MSS CRCs, where its presence correlates with immune exclusion and poor survival. Furthermore, we demonstrate that targeted depletion of canCBC cells enhances CD8⁺ T cell-mediated elimination of CRC organoids, providing functional evidence that canCBC cells actively contribute to immune resistance in MSS CRCs.

Our findings also show that canCBC cells are specifically associated with lymphatic dissemination, as their presence is enriched in CRCs with nodal involvement, suggesting a role in regional metastatic progression. This aligns with prior studies showing that tumor-initiating populations contribute to lymph node metastasis by facilitating immune evasion and enhancing invasive potential ^47,48^. Given that nodal metastasis is an early step in tumor dissemination, it will be important to determine whether distant metastatic lesions, such as those in the liver or lungs, also harbour an enrichment of canCBCs and potentially rely on the canCBC program for survival and immune resistance.

A key outstanding question is the precise mechanism by which canCBCs drive immune exclusion in MSS CRCs. One potential explanation is that their transcriptional program induces the secretion of immunosuppressive factors, such as NOTUM, which modulates WNT signalling and inhibits immune cell recruitment. Furthermore, our differential gene expression analysis suggests that canCBC cells could promote immune exclusion through a combined effect of multiple WNT/β-catenin signalling modulators, including NOTUM. Overall, these observations align with prior studies showing that dysregulated WNT signaling is a key driver of tumor immune escape and progression ^49,50^. Additionally, specific ablation of canCBC cells enhances CD8⁺ T cell-mediated tumor clearance, strongly suggesting that canCBC cells might directly suppress CD8⁺ T cell proliferation, inhibit their cytotoxic activity, or promote their exhaustion ^51^.

Epigenetic modifications may also play a role, as tumor-intrinsic epigenetic remodelling is known to suppress antigen presentation and impair CD8⁺ T cell recognition, a phenomenon observed in several immune-resistant tumors ^52–54^. Future studies integrating functional genomics and single-cell spatial transcriptomics will be essential to dissect the molecular pathways by which canCBCs orchestrate immune exclusion ^55,56^.

Another crucial question is whether canCBC cells can be therapeutically targeted to enhance immunotherapy responses. Given their stem-like properties and resilience to conventional treatments, as indicated by their role in promoting poor clinical outcome in CRC patients undergoing conventional chemoradiation-based therapeutics, strategies that disrupt their self-renewal capacity or modulate their immune interactions may improve treatment outcomes. While WNT pathway inhibitors have been explored in CRC, their broad inhibition leads to severe toxicity due to essential roles that the WNT pathway plays in maintaining normal tissue and immune homeostasis. A more selective approach, such as targeting canCBC program, which is specifically associated with the cancerous cells, may provide a therapeutic window to eliminate canCBC cells while preserving normal tissues, including routine epithelial turnover of the intestinal epithelium which relies on normal LGR5^+^ CBC cells ^9,30,31,57^.

From a clinical perspective, identifying canCBC cells as a distinct CSC subset provides a potential biomarker for MSS CRCs, particularly those predisposed to immune exclusion and therapy resistance. The feasibility of tissue-based profiling using single-cell transcriptomics or spatial multi-omics suggests that assessing canCBC prevalence could aid in patient stratification for immunotherapy. Furthermore, targeted therapeutic interventions against canCBCs could enable the development of novel combination strategies, including: (1) adaptive cellular therapies that selectively target surface proteins specifically expressed by canCBC cells, (2) small-molecule inhibitors that disrupt the canCBC program, and (3) combination regimens incorporating immune checkpoint blockade to enhance tumor clearance.

This study identifies canCBCs as a novel CSC subset in MSS CRCs, linking them to immune exclusion and resistance to immunotherapy. By elucidating their immunosuppressive potential, our findings open new therapeutic avenues for targeting immune-resistant tumors. Future research should focus on defining the molecular dependencies of canCBC cells, developing selective therapeutic strategies, and assessing their clinical translational potential to enhance immunotherapy efficacy in MSS CRC patients.

## Methods

### Ethics

This study complies with all relevant ethical regulations. Human CRC patient samples were collected through the ‘A Comprehensive Gastrointestinal and Hepatopancreaticobiliary Tumor Bank’ project and were approved by the Health Research Ethics Board of Alberta (HREBA) under ethics ID HREBA.CC-16-0769. The work related to mice was reviewed and approved by the Life and Environmental Sciences Animal Care Committee (LESACC) under protocols AC22-0068 and AC23-0117, in accordance with Canadian Council on Animal Care guidelines.

### Development of Human Organoid Cultures

Tissues were processed within 12 hours of resection from the patient. Tissue samples were placed in Advanced DMEM (Gibco) containing GlutaMAX (1X, Gibco), 100 U/ml penicillin (Gibco), 100 μg/ml streptomycin (Gibco), and 1X HEPES (Gibco). Samples were subsequently washed 5 times with 1X PBS (Gibco) and mechanically dissected using a razor blade in a 10cm petridish (Falcon). Accumax (innovative cell technologies) was added to the samples for enzymatic digestion for 30 minutes. Following 30 minutes the sample was spun down at 1240 rcf for 5 minutes. The supernatant was removed, and 5 mL of pre-warmed trypsin was added to the tissue for cell dissociation. The sample was incubated for 5 minutes at 37 °C. This process was repeated an additional time and then added with 10% FBS (Gibco) and the sample was spun down at 1240 rcf for 5 minutes.

The supernatant was removed, and the pellet was resuspended with Matrigel (VWR). This mixture was plated on 24 well plates (Corning) and incubated for 15 minutes in an incubator at 37 °C and 5% CO2 to allow time for the Matrigel to settle. Human organoid media was added to each well which comprised of Advanced DMEM (Gibco) containing GlutaMAX (1X, Gibco), 100 U/ml penicillin, 100 μg/ml streptomycin (Gibco), HEPES (1X Gibco), 1X B27 (invitrogen), 10% FBS (Gibco), 50 ng/mL mouse recombinant EGF (Invitrogen), 100ng/mL beta-EGF (Gibco), 10 µM Y27632 (Sigma), 1 µM NECA (Sigma), and 1 µM SB202190 (Sigma). Organoids were passaged every 7 says for maintenance.

### Formation of AKP organoids

Wildtype mouse organoids derived from male and female C57BL/6 mice were isolated from crypts as previously described ^58^. Constructs to induce Apc and Tp53 knockouts were done using plasmid #98293 and methods previously ^59^. New sgRNA for Apc and Tp53 were created and cloned into Addgene plasmid #98293. pHAGE-KRAS-G12D vector ^43^ (Addgene plasmid #116432) was used to induce a Kras Mutation in the wildtype intestinal organoids.

Wildtype organoids were mechanically broken down using a 12mL syringe and 16-gauge blunt needle. These organoids were then treated with 5 mL of pre-warmed trypsin and incubated for 5 minutes at 37 °C. This process was repeated an additional time, and 10% FBS (Gibco) was added to the sample and spun down at 1240 rcf for 5 minutes. Cells were mixed with a mixture of 295 µL of Opti-MEM (Gibco), 5 µL of lipofectamine 3000 (Invitrogen), 1 µg of each plasmid 1X B27 (invitrogen), 1X N2 (Invitrogen), 1mM of N-acetylcysteine (Sigma) and Advanced DMEM (Gibco) and plated on top of 80 µL of Matrigel (VWR) in a 24 well plate (Corning). Cells were left for 16 hours and subsequently media was removed and 60 µL of Matrigel (VWR) was added on top. Media containing 1X B27 (invitrogen), 1X N2 (Invitrogen), 1mM of N-acetylcysteine (Sigma) and Advanced DMEM (Gibco) was added, and organoids were allowed to grow. Knockouts and mutations were validated using qPCR.

This method was also conducted following the formation of AKP organoids for cOVA truncated protein using the Plasmid #25097. Positive transfections were selected by the addition of 400 µg/mL of G418 (Gibco).

### Quantitative real-time PCR

1 µg of RNA was extracted from tissue using the RNeasy Mini Kit (Qiagen) and was reverse transcribed using a reverse transcription kit (Applied Biosystems). Quantification was done using SYBR Green (Bio-Rad) and a CFX Connected Real-Time PCR Detection System (Bio-Rad). Data was quantified using the ΔΔCt method normalized to the expression of Actb

### Single-molecule FISH

Single-molecule fluorescence in situ hybridization (sm-FISH; RNAscope) was done on paraffin-embedded sections according to the manufacturer’s recommendations, using the RNAscope Multiplex Fluorescent Reagent Kit v2 Assay (ACD). Probes for LGR5, CD8A, and NOTUM were designed by the manufacture (ACD). Manufactures recommendations were followed and TSA dyes (ACD) 520, 570, and 650 were used for LGR5, CD8A, and NOTUM respectively. Samples were mounted and counterstained using Prolong Gold antifade with DAPI (manufacture) and imaged on a Nikon A1R+ confocal microscope.

### Single-cell RNA sequencing PDO sample preparation

PDOs were mechanical dissociated using a 12 mL syringe and 16-gauge needle followed by the addition of 5 mL of pre-warmed trypsin and incubated for 5 minutes at 37 °C. The addition of trypsin and incubation at 37 °C was repeated 3 additional times. 10% FBS (Gibco) was added and the sample and spun down at 1240 rcf for 5 minutes. Samples were washed twice with 1X PBS (Gibco) to remove residual trypsin. Cells were then stained with trypan blue (Gibco) and counted on a hemocytometer to access cell quantity and viability. Samples were run using the 3’ Chromium Single Cell 3’ Reagent Kit (10X genomics) and 10, 000 cell recovery was targeted.

### Single-cell RNA sequencing PDO Pre-processing

Fastq files were processed through Cell ranger v7.0.1 and sequenced were aligned against GRCh38. Recommended parameters were used. Cell ranger outputs were converted into Seurat objects using Seurat version 4.0.1 and corresponding metadata was added. Normal human organoid samples were added from GSE166556 and other colorectal cancer organoids were added from GSE121445. These samples were converted to Seurat objects, and all samples were merged into one object.

Quality control thresholds for mitochondria DNA percentage and the counts of transcripts was applied to each dataset. The top 2000 variable features were identified in this cohort of samples and were used to scale the data. PCA was used with 50 total principal components. The package Harmony was used, and the first 5 embeddings were used for further analysis. Production of the UMAP used 25 dimensions total and the resolution was set to 0.25. Identification of normal and cancer subtype clusters were done using genes highly expressed from previous human CRC single-cell analysis to identify revSC, canCBC, Cluster #6 and Normal CBC. These highly expressed genes were also visualized using a Dotplot to identify the percentage of cells expressing the gen and the level of expression.

### Single-cell RNA sequencing Human Sample Pre-processing

Samples from GSE178341, GSE201349, GSE132465, GSE166555, and GSE225857 were extracted and samples were converted to Seurat objects using Seurat version 4.0.1 and corresponding metadata was also added. All samples were merged together and filtered to have cells containing more than 400 counts and less that 25% mitochondria DNA. We selected the top 2000 variable genes, scaled the data and selected 50 principal components to run principal component analysis (PCA). To combat batch effect the Harmony package was used to make embeddings ^28^. Default Harmony parameters were used with 5 total embeddings merged. 1:15 dimensions were used when clustering the data for visualization using 0.25 resolution and Louvain algorithm.

Re-clustering with only epithelial cancer cells was done using the same parameters as previously described. Cluster types were identified using known cell markers for secretory, absorptive, proCSC, and revCSC. Cell signatures of these genes were also used to confirm the identities of these clusters.

### Single-cell and bulk RNA-seq interface and survival analysis

To integrate single-cell and bulk RNA-seq data, we applied Gene Set Variation Analysis (GSVA) to stratify 427 CRC patients from the TCGA-COAD dataset who had undergone chemoradiation therapy with clinical follow-up data. Patients were categorized into binary subcohorts based on CSC subtype enrichment, defining them as either enriched (‘+’) or deficient (‘–’) for each CSC-specific transcriptional program. Survival outcomes were assessed using the Kaplan-Meier estimator, with statistical significance determined by the log-rank test, which compared survival distributions between enriched and deficient patient subgroups. Hazard ratios and confidence intervals were estimated using Cox proportional hazards regression models to quantify the impact of CSC subtype enrichment on patient survival. This analysis revealed that patients with high revCSC or canCBC enrichment exhibited significantly worse survival compared to other CSC subtypes, reinforcing the clinical relevance of these populations in therapy resistance and disease progression.

### Cell cycle phase distribution

Cell cycle marker genes were obtained through previous publication that mark cells in G1, G2M, and S phase ^60^. These signatures were plotted on the human CRC scRNA-seq data and cells expressing an average log2 expression of the gene signature were deemed to be in one of the cell cycle phases. Cancer cell subtypes were subset individually and the number of cells in each phase of each subtype was converted to a percentage of the total number of cells of the cell type. These values were then compared in a bar graph to identify cell cycle differences for each cancer cell type.

### Calculating cancer cell type contribution in human samples

Following initial clustering of human cancer epithelial cells samples from GSE178341 were subset as each sample had microsatellite status in the metadata. The total number of cells from each sample was calculated with the number of cells that came from each cancer associated cell type. These values were converted to percentages to identify the percentage of cells contributing to each cancer associated cell type from each individual sample. These values were compared by MSS or MSI-H status within the metadata.

This analysis was also done on the total cohort of 176 samples comprised of 140 CRC patients in total from 5 different CRC patient cohorts. The percentage contribution for each cancer cell type was calculated and tumor cells versus normal cells was analysed. A paired student t-test was applied to identify any significance between cancerous and normal tissue samples.

### Differential gene expression analysis

A MAST test was used to identify differentially expressed genes unique to each cluster. Test were conducted for each cluster in comparison to all other clusters in the dataset. Minimum percentage, return thresholds, and logfc thresholds were all set to 25 percent.

### CD8a+ T Cell Isolation

Spleens were removed from OT1 mice aged from 8-24 weeks. The mouse was then sprayed with 70% ethanol until hairs were drenched. An incision was made in the abdominal area using surgical scissors to expose the peritoneal cavity and the spleen was removed aseptically. Spleens were first cut into three even sections and added to a mixture of 10 mL of RPMI (Gibco) and 10% FBS (Gibco) in a 10cm petridish (Falcon). The pieces of the spleen were mechanically crushed using the plunger of a 12 mL syringe. A 70 µm cell strainer (Falcon) was placed on a 50mL tube (Falcon) and 2mL of RPMI (Gibco) solution added to prime it. The resulting solution was filtered using a 70 μm filter (Falcon)to remove excess debris and the supernatant was centrifuged at 300 rcf for 5 minutes. The resulting pellet was resuspended in 5 mL ammonium chloride (Stem Cell Technologies), approximately 9x the volume of the pellet, and incubated in ice for 5 minutes to lyse the red blood cells in the pellet. 50 mL 1X PBS (Gibco) was added to the mixture and mixed using a pipette to neutralize the reaction. The solution was centrifuged at 500 rcf for 5 minutes, and the resulting pellet was resuspended in RPMI (Gibco). A portion of the resuspended pellet was mixed with trypan blue (Gibco) and counted on a hemocytometer to determine T cell quantity and viability.

Following this, CD8a+ cells were selectively isolated using the MidiMACS™ Separator and Starting Kit and the CD8a+ T Cell Isolation Kit (Miltenyi Biotec, #130-104-075 and #130-090-329 respectively), according to the manufacturer’s instructions.

### CD8a+ T Cell Activation

Isolated CD8+ T cells were directly plated on 6 well plates (Falcon) pre-coated with anti-CD3 and anti-CD28 (BD Bioscience). CD8+ cells were added with 4.45 mL RPMI (Gibco), 10% FBS (Gibco), 50 µM 2-mercaptoethanol (Gibco), 1X Insulin-Transferrin-Selenium (Gibco), 1X L-Glutamine (Gibco), 100 U/ml penicillin (Gibco), 100 μg/ml streptomycin (Gibco), 1 ng/mL IL-7 (Invitrogen), and 2.5 ng/mL IL-2 (Gibco). These cultures were incubated at 37 °C and 5% CO2 for two days before being used directly in organoid-CD8+ T cell co-culture assays.

### Apcdd1 ablation vector construct

This protocol was adapted from Shimokawa and colleagues ^5^. Two segments of the Apcdd1 gene (both ∼1 kbp) were PCR amplified to replace Lgr5 arms from the HR180-LGR5-iCT vector (Addgene plasmid #129094). Briefly, the first segment, termed the 5’ homology arm, ended 27 nucleotides upstream of the gene translation start site and extended upstream. The second segment, termed the 3’ homology arm, started exactly at the translation start site and extended downstream. Each of these homology arms were amplified using C57 mouse DNA extracted from mouse tail snips using a DNA extraction kit (Extracta DNA Prep for PCR, Quantabio). 1 µg of DNA was used with 1 µM of forward and reverse primers with 0.5 µL of High-Fidelity Taq polymerase (Takara Bio) and water topped up to 50 µL. PCR cycles were run using manufacture recommendations.

1 µg 5’ homology arm was digested along with the Plasmid #129094 with 1µL enzymes NdeI (NEB), 1 µL enzyme SacI-HF (NEB), 5 µl of custmart buffer (NEB) topped up with water to 50 µL total reaction. Reactions were left in 37 °C for one hour to complete. 1 µg 3’’ homology arm was digested along with the Plasmid #129094 with 1µL enzymes BsaBI (NEB), 1 µL enzyme SphI-HF (NEB), 5 µl of custmart buffer (NEB) topped up with water to 50 µL total reaction. Reactions were left in 37 °C for 30 minutes and then 60 °C for 30 minutes for the reaction to complete. This would create overhangs for ligation of the arms into the Plasmid #129094.Ligation of each of the arms was done using 50ng of the cut 5’ and 3’ sections of Plasmid #129094 with 12.5 ng of the digest homology arm, 5 µL of 2X Quick Ligase Buffer (NEB), 1 µL of Quick Ligase (NEB) topped with water for a 11 µL total reaction. The reaction was incubated at room temperature for 10 minutes followed by 16 °C for one hour. Ligation product was transformed into StbI3 bacteria (Invitrogen) and manufacture recommendations were followed. CRISPR-Cas9 transfer vector was designed by cloning sgRNA targeting the region between the 5’ and 3’ homology arms of the Apcdd1 into the lentiCRISPRv2 hygro plasmid (Addgene plasmid #98291). This vector was transfected alongside the Apcdd1 ablation vector to facilitate the incorporation of the ablation vector through homologous recombination. Selection was done using 4 µg/mL of puromycin (ThermoFisher) for 48 hours. Ablation was induced by the addition of 4nM of AP20187 (Sigma) for 48 hours.

### Organoid Fixing and Staining

AKP cOVA organoids were plated in 12 µL of Matrigel (VWR) on 10mm circular coverslips (ThermoFisher) and grown in AKP organoid media as stated earlier. Samples were allowed to grow for 3 days. 1 mL of PBST was added to each well with the addition of two drops of NucRed Dead 647 (Invitrogen) and incubated for 30 minutes. Samples were then fixed in 2 mL of formalin (Sigma) for 45 minutes. Organoids were washed 3 times with a mixture of 1X PBS (Gibco) and 0.2% Tween (Sigma) (PBST). Following 30 minutes wells were washed twice with 1 mL PBST and 20 µM of Hoechst (ThermoFisher) was added to fresh PBST and incubated for 30 minutes. Wells were washed twice with 1 mL PBST and circular coverslips (ThermFisher) were mounted with 10 µL ProLong Gold Antifade (Invitrogen).

### Organoid-CD8+ T Cell Coculture

CD8a+ T cells cultured for 2 days, as described above, were scraped off the bottom of the well to lift T cells. T cells and T cell media was transferred to a 15 mL tube (Falcon) and centrifuged at 500 rcf for 5 minutes. Supernatant was then removed, and pellet was resuspended in 4.45 mL RPMI (Gibco), 10% FBS (Gibco), 50 µM 2-mercaptoethanol (Gibco), 1X Insulin-Transferrin-Selenium (Gibco), 1X L-Glutamine (Gibco), 100 U/ml penicillin (Gibco), 100 μg/ml streptomycin (Gibco), 1 ng/mL IL-7 (Invitrogen), and 2.5 ng/mL IL-2 (Gibco). In parallel, organoids were passed through a 12 mL syringe (ThermoFisher) and blunt needle (ThermoFisher) and plated in 35 µL of Matrigel (VWR) on a 24 well plate. These cells were given organoid media as stated earlier and allowed to grow for 48 hours. Following organoids were put into a 15 mL tube (Falcon) and spun down at 1240 rcf. The supernatant was taken out and the pellet was resuspended in 20 µL Matrigel (VWR) and 500, 000 OT-1 CD8a+ T cells. This mixture was then platted on 10 mm circular coverslips (ThermoFisher) and let to set for 15 minutes at 37 °C. 1 mL of T cell medium as described before was added to the co-culture mixture and samples were cultured for 48 hours at 37 °C and 5% CO2. Following 48 hours samples were fixed and staining procedure was conducted. 2

## Acknowledgements

This study was supported by an Arnie Charbonneau Cancer Institute’s Team Accelerator Grant awarded to A.A. J.C. and P.S. were supported by student awards from NSERC and Biotherapeutics for Cancer Treatment (BioCanRx), respectively.

---

**Extended Data Figure 1.**
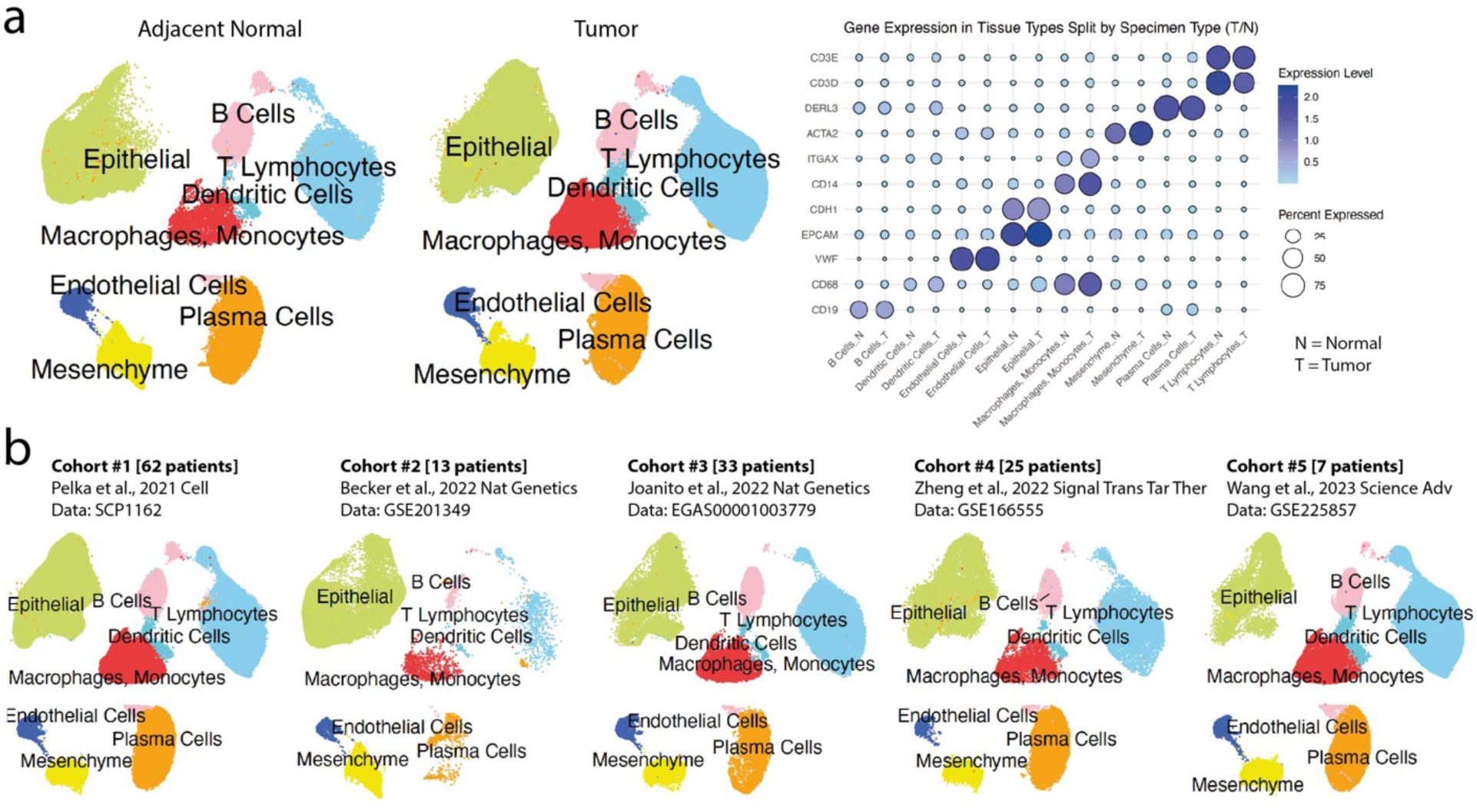
Identification of Distinct Transcriptional Phenotypes Across Five CRC Patient Cohorts. **a**, A UMAP visualization of single-cell transcriptomes of cancerous cells and normal cells, along with the associated tissues, from five independent CRC patient cohorts, along with expression of lineage specific markers shown in the form of self-explanatory dotplot. **b**, Individual UMAPs representing a single CRC patient cohort, across five independently generated datasets by multiple research teams.

**Extended Data Figure 2.**
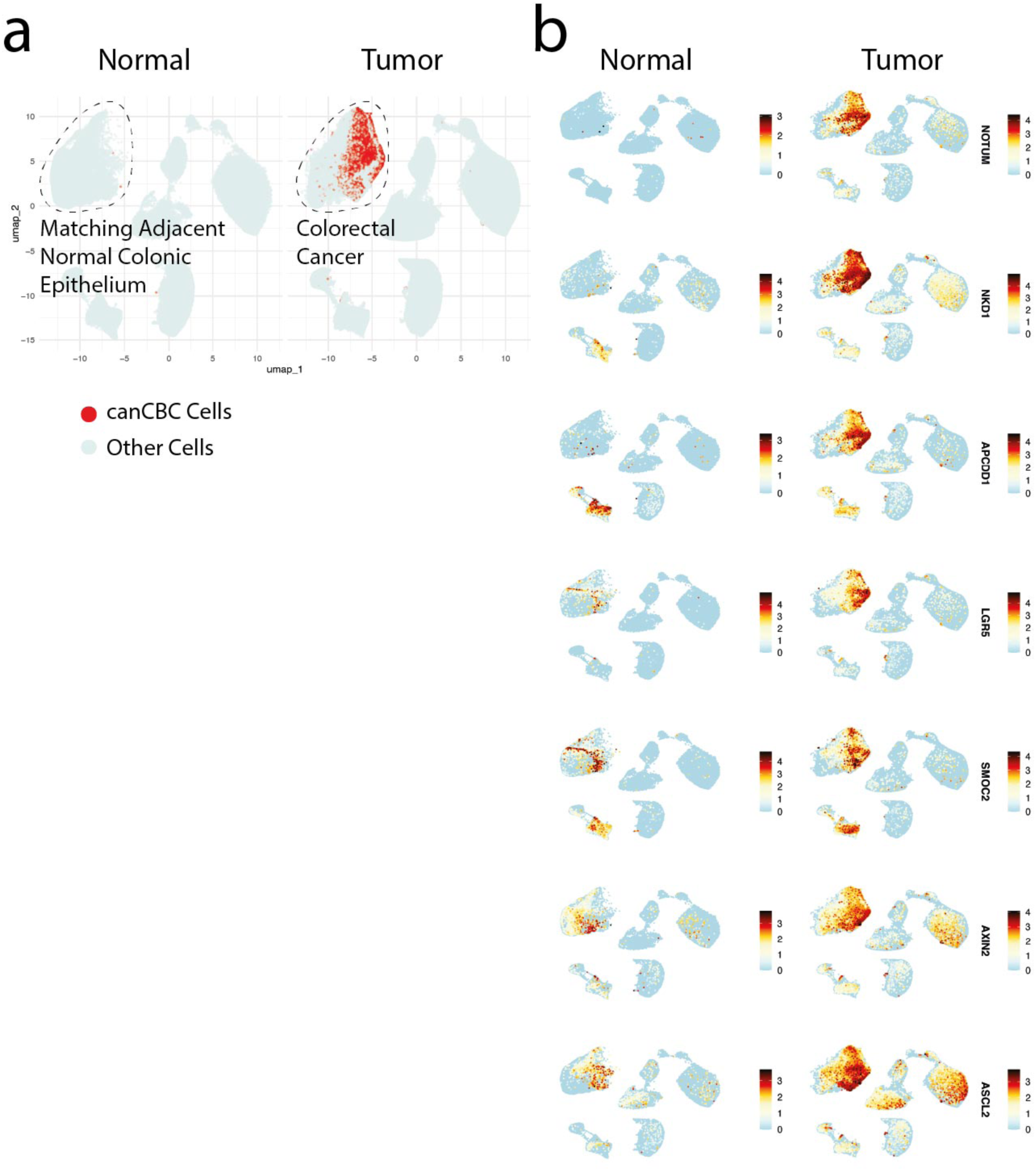
Gene Expression Analysis in Cancerous and Normal Tissues. **a**, Distribution of canCBC cells, defined by the average expression of canCBC marker genes NOTUM, APCDD1, and NKD1. **b**, UMAP visualization of gene expression levels for the indicated genes in cancerous and normal tissues.

**Extended Data Figure 3.**
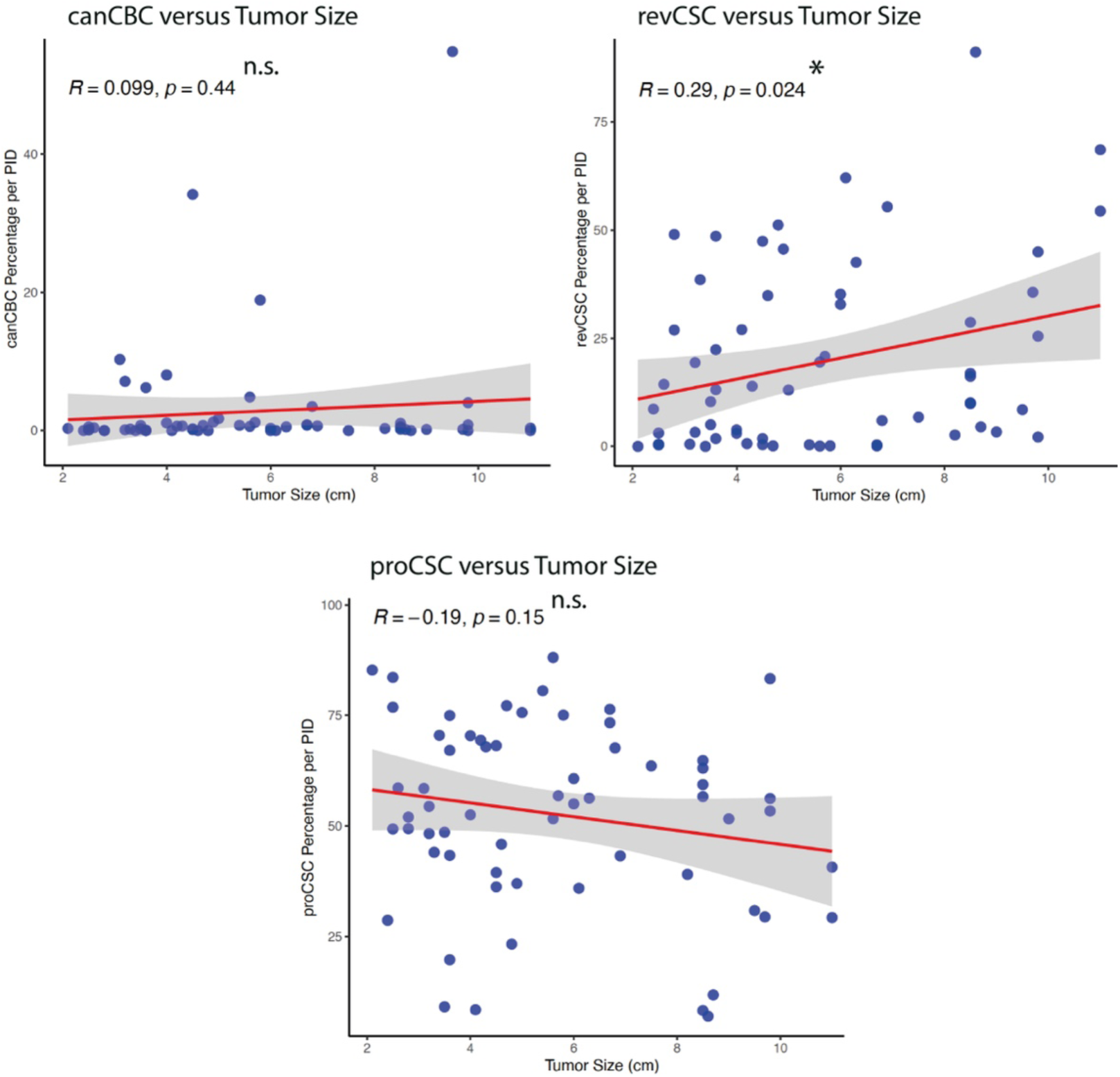
Tumor Size and Enrichment of canCBC Program. A correlation plot showing the relationship between tumor size and the percentage of canCBCs, revCSCs and proCSCs per patient (PID). Pearson correlation was used to assess statistical significance, showing no significant correlation. A trendline (linear regression) is included for visualization. Each point represents an individual patient, with the x-axis indicating tumor size (cm) and the y-axis representing the percentage of canCBC cells relative to total tumor cells per PID. n.s., non-significant.

**Extended Data Figure 4.**
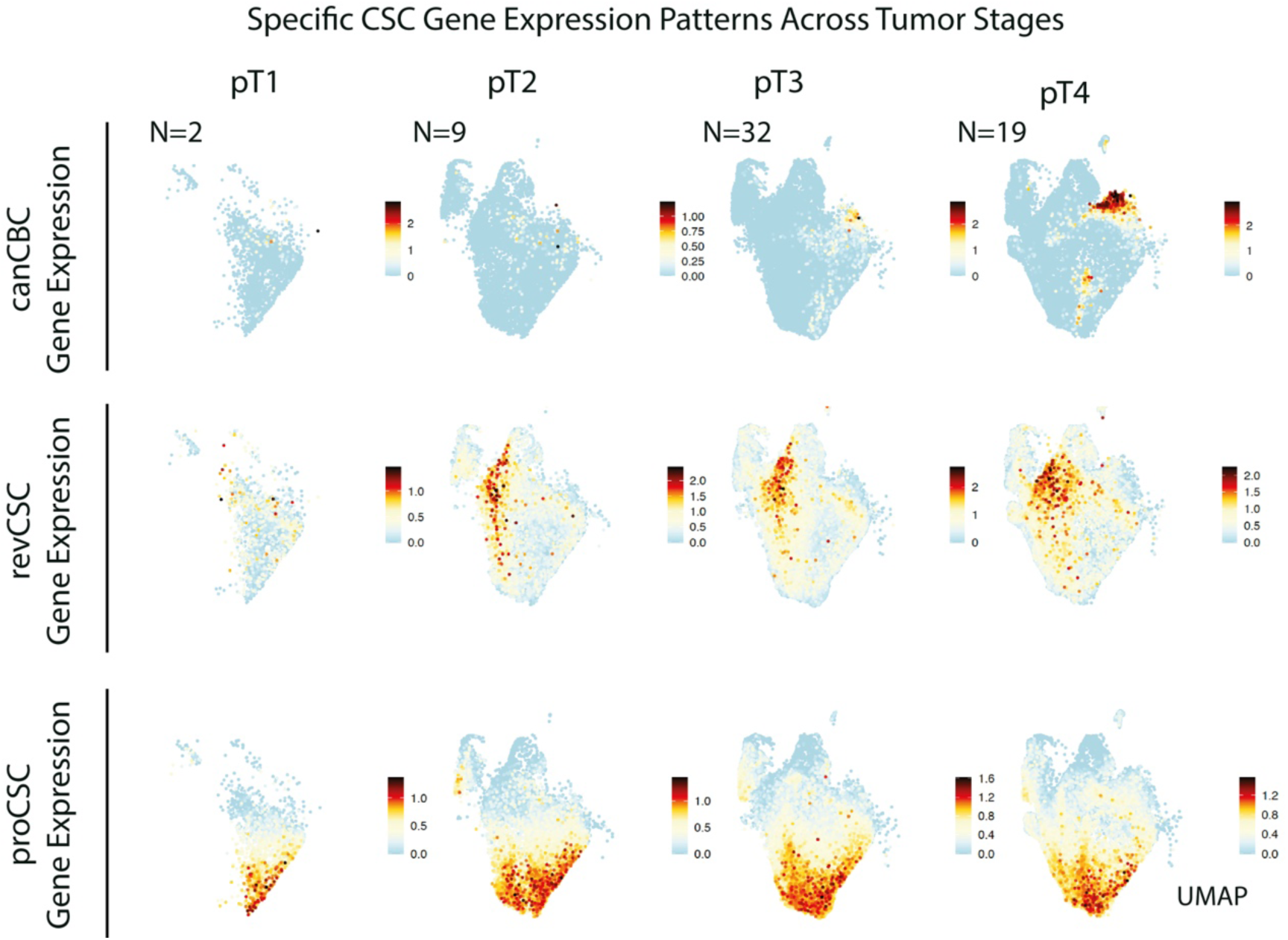
canCBC Association with Tumor Stages. A UMAP visualization showing the average expression of canCBC, revCSC and proCSC signature genes across all four tumor stage categories. ’N’ represents the number of CRC patients.

**Extended Data Figure 5.**
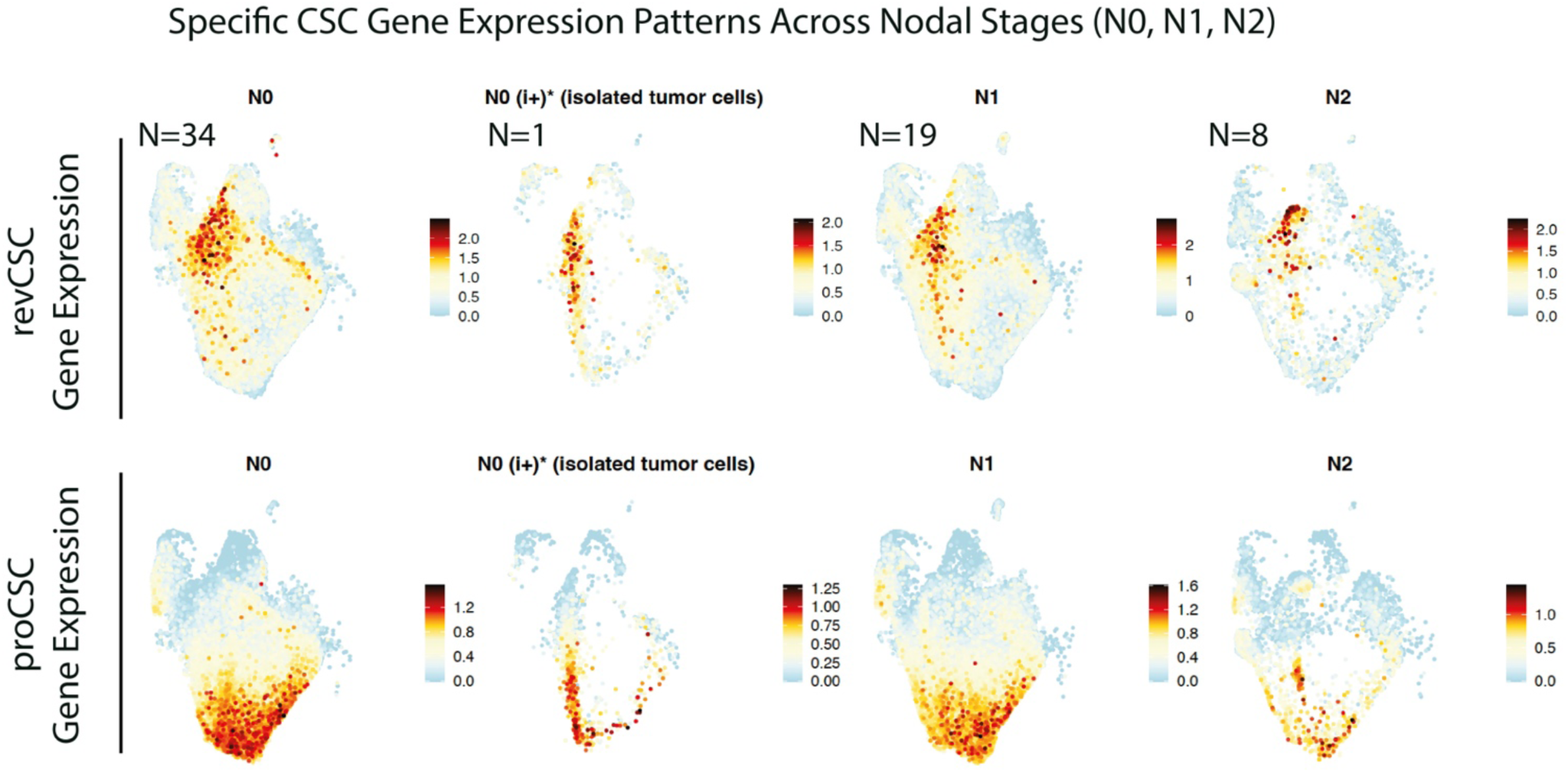
Association of revCSCs and proCSCs with Nodal Stages. A UMAP visualization showing the average expression of revCSC and proCSC signature genes across nodal categories. ’N’ represents the number of CRC patients.

**Extended Data Figure 6.**
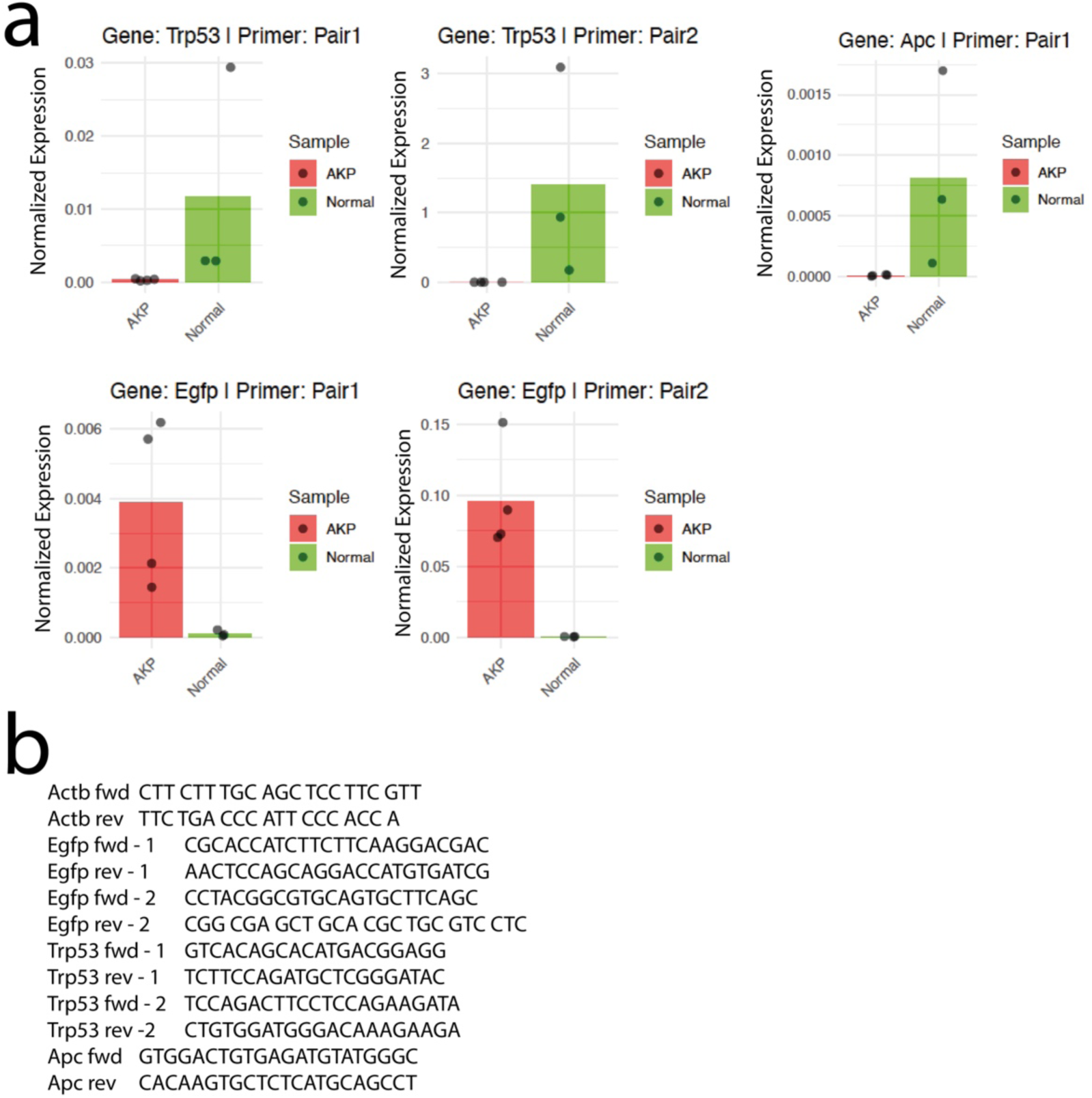
Validating AKP Tumor Organoid Genotype. **a**, Quantitative RT-PCR results for the indicated genes in AKP (cancer) and normal organoids, with expression values normalized to the housekeeping gene Actb. **b**, Primer sequences used in the experiment are provided.

**Extended Data Figure 7.**
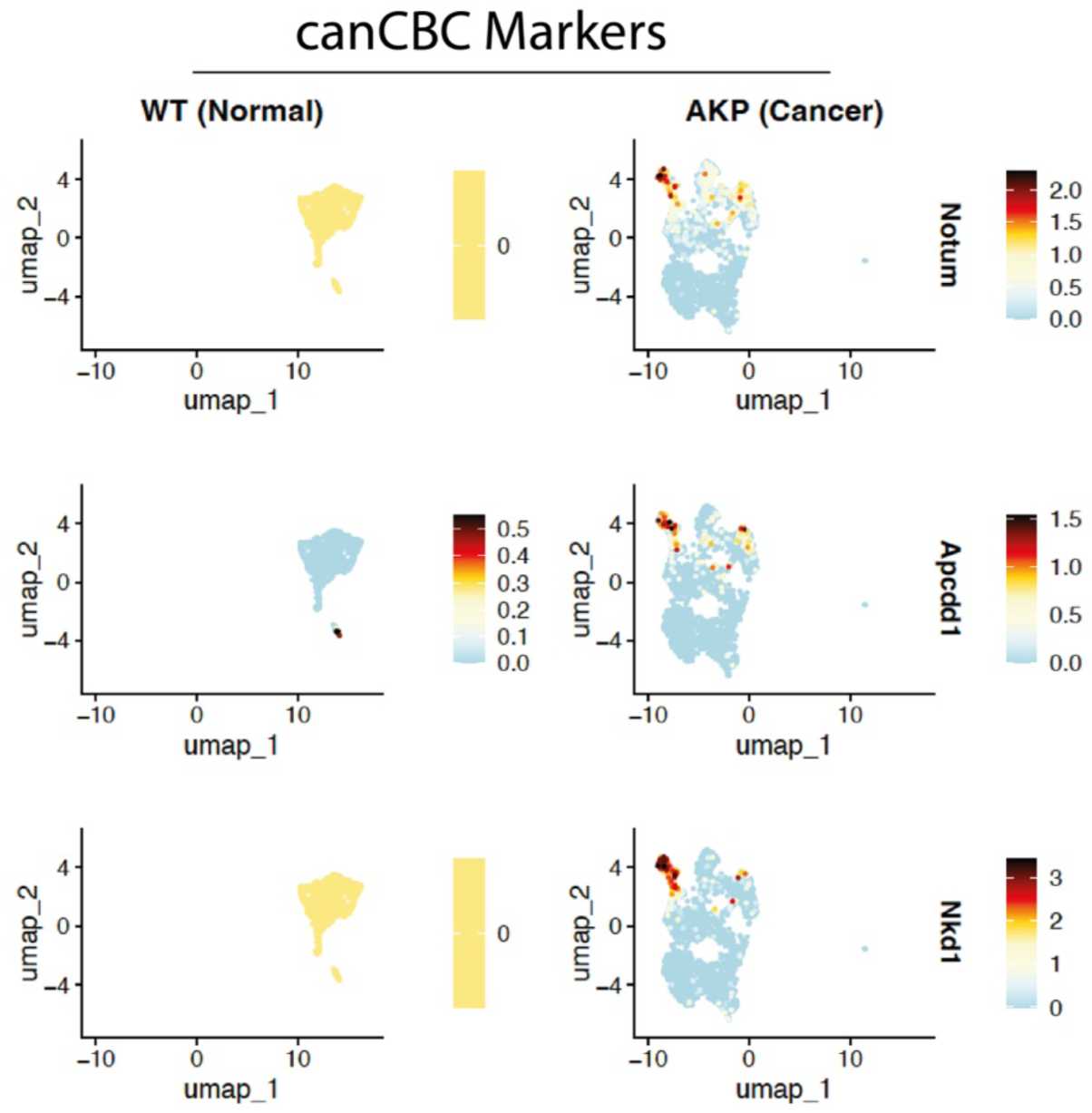
Genes Uniquely Expressed in canCBC Cells. UMAP visualization of gene expression levels for Notum, Apcdd1, and Nkd1. Expression of all three genes is restricted to a single cell cluster that represents a relatively rare population within AKP mouse organoids.

**Extended Data Figure 8.**
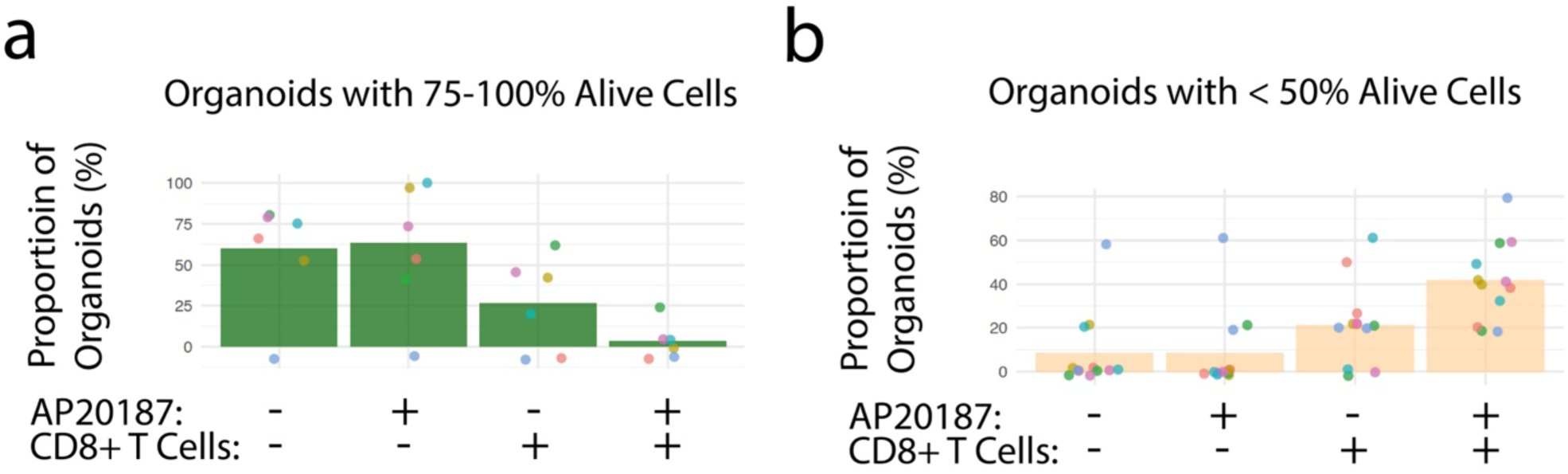
Co-cultures of CD8+ T Cells and canCBC-ablated AKP Organoids Facilitates Efficient Tumor Removal. **a**,**b**, Number of AKP tumor organoids that contain either more than 75% viable cells (**a**) or less than 50% viable cells (**b**) detected upon indicated treatments are shown.

## References

1. Siegel, R. L., Miller, K. D., Wagle, N. S. & Jemal, A. Cancer statistics, 2023. CA Cancer J Clin 73, 17–48 (2023).

2. Horn, L. A., Fousek, K. & Palena, C. Tumor Plasticity and Resistance to Immunotherapy. Trends in Cancer Preprint at 10.1016/j.trecan.2020.02.001 (2020).

3. Shin, A. E., Giancotti, F. G. & Rustgi, A. K. Metastatic colorectal cancer: mechanisms and emerging therapeutics. Trends Pharmacol Sci 44, 222 (2023).

4. Dienstmann, R. et al. Consensus molecular subtypes and the evolution of precision medicine in colorectal cancer. Nat Rev Cancer 17, 79–92 (2017).

5. Shimokawa, M. et al. Visualization and targeting of LGR5 + human colon cancer stem cells. Nature 545, 187–192 (2017).

6. De Sousa E Melo, F., et al. A distinct role for Lgr5 + stem cells in primary and metastatic colon cancer. Nature (2017) doi:10.1038/nature21713.

7. Leushacke, M. et al. Lgr5-expressing chief cells drive epithelial regeneration and cancer in the oxyntic stomach. Nat Cell Biol 19, 774–786 (2017).

8. Lenos, K. J. et al. Molecular characterization of colorectal cancer related peritoneal metastatic disease. Nature Communications 2022 13:1 13, 1–14 (2022).

9. Ayyaz, A. et al. Single-cell transcriptomes of the regenerating intestine reveal a revival stem cell. Nature 569, 121–125 (2019).

10. Fink, M. et al. Chromatin remodelling in damaged intestinal crypts orchestrates redundant TGFβ and Hippo signalling to drive regeneration. Nature Cell Biology 2024 1–15 (2024) doi:10.1038/s41556-024-01550-4.

11. Merlos-Suárez, A. et al. The intestinal stem cell signature identifies colorectal cancer stem cells and predicts disease relapse. Cell Stem Cell 8, 511–524 (2011).

12. Tape, C. J. Plastic persisters: revival stem cells in colorectal cancer. (2024) doi:10.1016/j.trecan.2023.11.003.

13. Qin, X. et al. An oncogenic phenoscape of colonic stem cell polarization. Cell 186, 5554–5568.e18 (2023).

14. Le, D. T. et al. Mismatch repair deficiency predicts response of solid tumors to PD-1 blockade. Science 357, 409–413 (2017).

15. Overman, M. J. et al. Durable clinical benefit with nivolumab plus ipilimumab in DNA mismatch repair-deficient/microsatellite instability-high metastatic colorectal cancer. Journal of Clinical Oncology 36, 773–779 (2018).

16. Llosa, N. J. et al. The vigorous immune microenvironment of microsatellite instable colon cancer is balanced by multiple counter-inhibitory checkpoints. Cancer Discov 5, 43–51 (2015).

17. Cristescu, R. et al. Pan-tumor genomic biomarkers for PD-1 checkpoint blockade-based immunotherapy. Science 362, (2018).

18. Postow, M. A., Sidlow, R. & Hellmann, M. D. Immune-Related Adverse Events Associated with Immune Checkpoint Blockade. New England Journal of Medicine 378, 158–168 (2018).

19. Martins, F. et al. Adverse effects of immune-checkpoint inhibitors: epidemiology, management and surveillance. Nature Reviews Clinical Oncology 2019 16:9 16, 563–580 (2019).

20. Agudo, J. et al. Quiescent Tissue Stem Cells Evade Immune Surveillance. Immunity (2018) doi:10.1016/j.immuni.2018.02.001.

21. Miao, Y. et al. Adaptive Immune Resistance Emerges from Tumor-Initiating Stem Cells. Cell (2019) doi:10.1016/j.cell.2019.03.025.

22. Jia, L., Zhang, W. & Wang, C. Y. BMI1 Inhibition Eliminates Residual Cancer Stem Cells after PD1 Blockade and Activates Antitumor Immunity to Prevent Metastasis and Relapse. Cell Stem Cell (2020) doi:10.1016/j.stem.2020.06.022.

23. Pelka, K. et al. Spatially organized multicellular immune hubs in human colorectal cancer. Cell 184, 4734–4752.e20 (2021).

24. Becker, W. R. et al. Single-cell analyses define a continuum of cell state and composition changes in the malignant transformation of polyps to colorectal cancer. Nature Genetics 2022 54:7 54, 985–995 (2022).

25. Lee, H. O. et al. Lineage-dependent gene expression programs influence the immune landscape of colorectal cancer. Nature Genetics 2020 52:6 52, 594–603 (2020).

26. Uhlitz, F. et al. Mitogen-activated protein kinase activity drives cell trajectories in colorectal cancer. EMBO Mol Med 13, (2021).

27. Wang, F. et al. Single-cell and spatial transcriptome analysis reveals the cellular heterogeneity of liver metastatic colorectal cancer. Sci Adv 9, (2023).

28. Korsunsky, I. et al. Fast, sensitive, and accurate integration of single cell data with Harmony. Nat Methods 16, 1289 (2019).

29. Bala, P. et al. Aberrant cell state plasticity mediated by developmental reprogramming precedes colorectal cancer initiation. Sci Adv 9, (2023).

30. Flanagan, D. J. et al. NOTUM from Apc-mutant cells biases clonal competition to initiate cancer. Nature 2021 594:7863 594, 430–435 (2021).

31. van Neerven, S. M. et al. Apc-mutant cells act as supercompetitors in intestinal tumour initiation. Nature 2021 594:7863 594, 436–441 (2021).

32. Wang, R. et al. Systematic evaluation of colorectal cancer organoid system by single-cell RNA-Seq analysis. Genome Biol 23, (2022).

33. Joanito, I. et al. Single-cell and bulk transcriptome sequencing identifies two epithelial tumor cell states and refines the consensus molecular classification of colorectal cancer. Nat Genet 54, 963–975 (2022).

34. van de Moosdijk, A. A. A., van de Grift, Y. B. C., de Man, S. M. A., Zeeman, A. L. & van Amerongen, R. A novel Axin2 knock-in mouse model for visualization and lineage tracing of WNT/CTNNB1 responsive cells. Genesis 58, e23387 (2020).

35. Li, J., Fu, Y., Zhang, K. & Li, Y. Integration of Bulk and Single-Cell RNA-Seq Data to Construct a Prognostic Model of Membrane Tension-Related Genes for Colon Cancer. Vaccines (Basel) 10, (2022).

36. Yalcin-Ozkat, G. Molecular Modeling Strategies of Cancer Multidrug Resistance. Drug Resist Updat 100789 (2021) doi:10.1016/J.DRUP.2021.100789.

37. Thomas, E. M. et al. Advancing translational research for colorectal immuno-oncology. British Journal of Cancer 2023 129:9 129, 1442–1450 (2023).

38. Ganesh, K. et al. Immunotherapy in colorectal cancer: rationale, challenges and potential. Nature Reviews Gastroenterology & Hepatology 2019 16:6 16, 361–375 (2019).

39. Sharma, P., Hu-Lieskovan, S., Wargo, J. A. & Ribas, A. Primary, Adaptive, and Acquired Resistance to Cancer Immunotherapy. Cell (2017) doi:10.1016/j.cell.2017.01.017.

40. André, T. et al. Pembrolizumab in Microsatellite-Instability–High Advanced Colorectal Cancer. New England Journal of Medicine 383, 2207–2218 (2020).

41. Zhang, Y. et al. Impact of lymph node metastasis on immune microenvironment and prognosis in colorectal cancer liver metastasis: insights from multiomics profiling. British Journal of Cancer 2025 1–12 (2025) doi:10.1038/s41416-024-02921-2.

42. Hogquist, K. A. et al. T cell receptor antagonist peptides induce positive selection. Cell 76, 17– 27 (1994).

43. Ng, P. K. S. et al. Systematic Functional Annotation of Somatic Mutations in Cancer. Cancer Cell 33, 450–462.e10 (2018).

44. Liao, W. et al. KRAS-IRF2 axis drives immune suppression and immune therapy resistance in colorectal cancer. Cancer Cell 35, 559 (2019).

45. Tian, Y. et al. APC and P53 mutations synergise to create a therapeutic vulnerability to NOTUM inhibition in advanced colorectal cancer. Gut 72, 2294–2306 (2023).

46. Kreso, A. & Dick, J. E. Evolution of the cancer stem cell model. Cell Stem Cell 14, 275–291 (2014).

47. Chen, P., Hsu, W. H., Han, J., Xia, Y. & DePinho, R. A. Cancer Stemness Meets Immunity: From Mechanism to Therapy. Cell Rep 34, 108597 (2021).

48. Li, L. et al. Cancer stem cells promote lymph nodes metastasis of breast cancer by reprogramming tumor microenvironment. Transl Oncol 35, 101733 (2023).

49. Zhan, T., Rindtorff, N. & Boutros, M. Wnt signaling in cancer. Oncogene 36, 1461–1473 (2017).

50. Galluzzi, L., Spranger, S., Fuchs, E. & López-Soto, A. WNT Signaling in Cancer Immunosurveillance. Trends Cell Biol 29, 44–65 (2019).

51. Luke, J. J., Bao, R., Sweis, R. F., Spranger, S. & Gajewski, T. F. WNT/β-catenin Pathway Activation Correlates with Immune Exclusion across Human Cancers. Clin Cancer Res 25, 3074–3083 (2019).

52. Parab, A., Kumar Bhatt, L. & Omri, A. Targeting Epigenetic Mechanisms: A Boon for Cancer Immunotherapy. Biomedicines 11, 169 (2023).

53. Yang, J. et al. Epigenetic regulation in the tumor microenvironment: molecular mechanisms and therapeutic targets. Signal Transduction and Targeted Therapy 2023 8:1 8, 1–26 (2023).

54. Villanueva, L., Álvarez-Errico, D. & Esteller, M. The Contribution of Epigenetics to Cancer Immunotherapy. Trends Immunol 41, 676–691 (2020).

55. Spranger, S. & Gajewski, T. F. Impact of oncogenic pathways on evasion of antitumour immune responses. Nat Rev Cancer 18, 139–147 (2018).

56. Tian, X. Q. et al. Epigenetic silencing of LRRC3B in colorectal cancer. Scand J Gastroenterol 44, 79–84 (2009).

57. Barker, N. et al. Identification of stem cells in small intestine and colon by marker gene Lgr5. Nature 449, 1003–1007 (2007).

58. Sato, T. et al. Single Lgr5 stem cells build crypt-villus structures in vitro without a mesenchymal niche. Nature (2009) doi:10.1038/nature07935.

59. Doench, J. G. et al. Optimized sgRNA design to maximize activity and minimize off-target effects of CRISPR-Cas9. Nat Biotechnol (2016) doi:10.1038/nbt.3437.

60. Tirosh, I. et al. Dissecting the multicellular ecosystem of metastatic melanoma by single-cell RNA-seq. Science(1979) 352, 189–196 (2016).

